# Single-cell RNA-sequencing reveals pervasive but highly cell type-specific genetic ancestry effects on the response to viral infection

**DOI:** 10.1101/2020.12.21.423830

**Authors:** Haley E Randolph, Zepeng Mu, Jessica K Fiege, Beth K Thielen, Jean-Christophe Grenier, Mari S Cobb, Julie G Hussin, Yang I Li, Ryan A Langlois, Luis B Barreiro

**Affiliations:** Committee on Genetics, Genomics, and Systems Biology, University of Chicago, Chicago, IL, USA; Center for Immunology, University of Minnesota, Minneapolis, MN, USA; Department of Microbiology and Immunology, University of Minnesota, Minneapolis, MN, USA; Department of Pediatrics, Division of Pediatric Infectious Diseases and Immunology, University of Minnesota, Minneapolis, MN, USA; Research Center, Montreal Heart Institute, Montreal, QC, Canada; Section of Genetic Medicine, Department of Medicine, University of Chicago, Chicago, IL, USA; Faculty of Medicine, Université de Montréal, Montreal, QC, Canada; Department of Human Genetics, University of Chicago, Chicago, IL, USA; Committee on Immunology, University of Chicago, Chicago, IL, USA

**Keywords:** Genetic ancestry, single-cell RNA-sequencing, immune responses, influenza infection, natural selection, expression quantitative trait loci (eQTL)

## Abstract

Humans vary in their susceptibility to infectious disease, partly due to variation in the immune response following infection. Here, we used single-cell RNA-sequencing to quantify genetic contributions to this variation in peripheral blood mononuclear cells, focusing specifically on the transcriptional response to influenza infection. We find that monocytes are the most responsive to influenza infection, but that all cell types mount a conserved interferon response, which is stronger in individuals with increased European ancestry. By comparing European American and African American individuals, we show that genetic ancestry effects on expression are common, influencing 29% of genes, but highly cell type-specific. Further, we demonstrate that much of this population-associated expression variation is explained by *cis* expression quantitative trait loci, which are enriched for signatures of recent positive selection. Our findings establish common *cis*-regulatory variants—including those that are differentiated by genetic ancestry—as important determinants of the antiviral immune response.

## Introduction

Pathogenic viruses constitute one of the strongest sources of selective pressure in human evolutionary history (Barreiro et al., 2009; Enard and Petrov, 2018; Enard et al., 2016; Fumagalli et al., 2011; Siddle and Quintana-Murci, 2014). Prior to the modern era, however, global pandemics on the scale of the 1918 Spanish influenza or the ongoing SARS-CoV-2 pandemic were probably rare. Due to the restricted potential for long-distance (especially intercontinental) exchange, earlier viral epidemics are thought to have been strongly stratified by geography (Enard and Petrov, 2020). Consequently, although most human genetic variation is shared between populations, differences in viral-mediated selection pressures may have driven divergence in the frequencies of polymorphisms that mediate the viral host response (either because of differences in the viruses that caused epidemic outbreaks or heterogeneity in the timing of epidemic events between populations). If so, the pattern of past epidemic outbreaks may have contributed to variation in viral susceptibility observed within and among modern-day human populations— perhaps interacting with or compounding known health disparities that contribute to substantially higher rates of influenza and COVID-19 hospitalization in Black versus non-Hispanic white Americans (e.g., the Centers for Disease Control and Prevention (CDC) estimates a 79% higher rate of influenza-related hospitalizations for Black versus white Americans: (CDC, 2020)).

Genetics is thought to play an important role in explaining population variation in susceptibility to influenza and other viral pathogens (Albright et al., 2008; Kenney et al., 2017). Supporting this view, a study exploring the impact of regulatory genetic variation on gene expression levels (i.e. expression quantitative trait loci [eQTL] studies) following influenza A virus (IAV) infection of human dendritic cells revealed 121 genetic variants that are significantly associated with the immune response to IAV, including one *cis*-acting variant associated with the interferon regulatory factor 7 gene (*IRF7)* that also acts as a *trans*-regulator responsible for genetic effects on a hub of IRF7-induced antiviral genes (Lee et al., 2014). Variation in the gene expression response to IAV *in vitro* is also correlated with genetic ancestry in monocytes derived from individuals of African and European descent (Quach et al., 2016).

A major limitation of studies to date, however, is their focus on a single, isolated immune cell type. This approach is blind to genetic effects that act in a cell type-specific manner. Further, it fails to capture critical interactions between the array of immune cell types needed to mount an efficient response to a viral infection. To address these limitations, here, we combine single-cell RNA-sequencing with *in vitro* infection assays of peripheral blood mononuclear cells (PBMCs) with pathogenic influenza A virus. We identify both shared and cell type-specific responses to IAV that are detectable only when gene expression estimates are resolved into individual cell types. We then investigate the degree to which the transcriptional response to IAV is structured by European versus African genetic ancestry and dissect likely genetic contributions to these differences. Finally, we investigate whether variants that are associated with ancestry-related differences are likely to have evolved in response to past selection pressure. Our results show that *cis*-regulatory genetic variation contributes to phenotypic differences in the immune response of modern humans to IAV, including both within and between-population variation. Further, some of these variants—especially those linked to autoimmune risk—carry signatures of recent positive selection, particularly in Europeans, suggesting that at least some present-day autoimmune risk loci were adaptive and conferred a functional benefit during our evolutionary history.

### Single-cell profiling of the transcriptional response to influenza infection

We exposed peripheral blood mononuclear cells (PBMCs) sampled from a diverse panel of humans to either a mock treatment (negative control) or to the pandemic H1N1 Cal/04/09 influenza A virus (IAV) strain (multiplicity of infection [MOI] 0.5) (n = 180 samples, comprised of paired mock-exposed and IAV-infected samples from each of 90 individuals). Following 6 hours of exposure, we performed single-cell RNA-sequencing on all samples (Fig. 1A). In total, we captured 255,731 single-cell transcriptomes across all individuals and conditions (n = 235,161 high-quality cells retained after filtering, Table S1). In addition, we collected low-pass whole-genome sequencing for the same individuals (n = 89), which we used to estimate the proportion of African and European admixture for each individual. Self-identified African American (AA, n = 45) individuals had a modest, although highly variable, percentage of European ancestry (mean = 11%, range = 0 – 43%), while self-identified European American (EA, n = 44) individuals displayed more limited levels of African ancestry (mean = 1%, range = 0 – 23%) (Fig. S1A, Table S1). UMAP clustering revealed eight distinct immune cell types (Fig. 1B), with five major cell clusters corresponding to the five main cell types found in PBMCs, including CD4^+^ T cells, CD8^+^ T cells, B cells, natural killer (NK) cells, and monocytes.

**Fig. 1.**
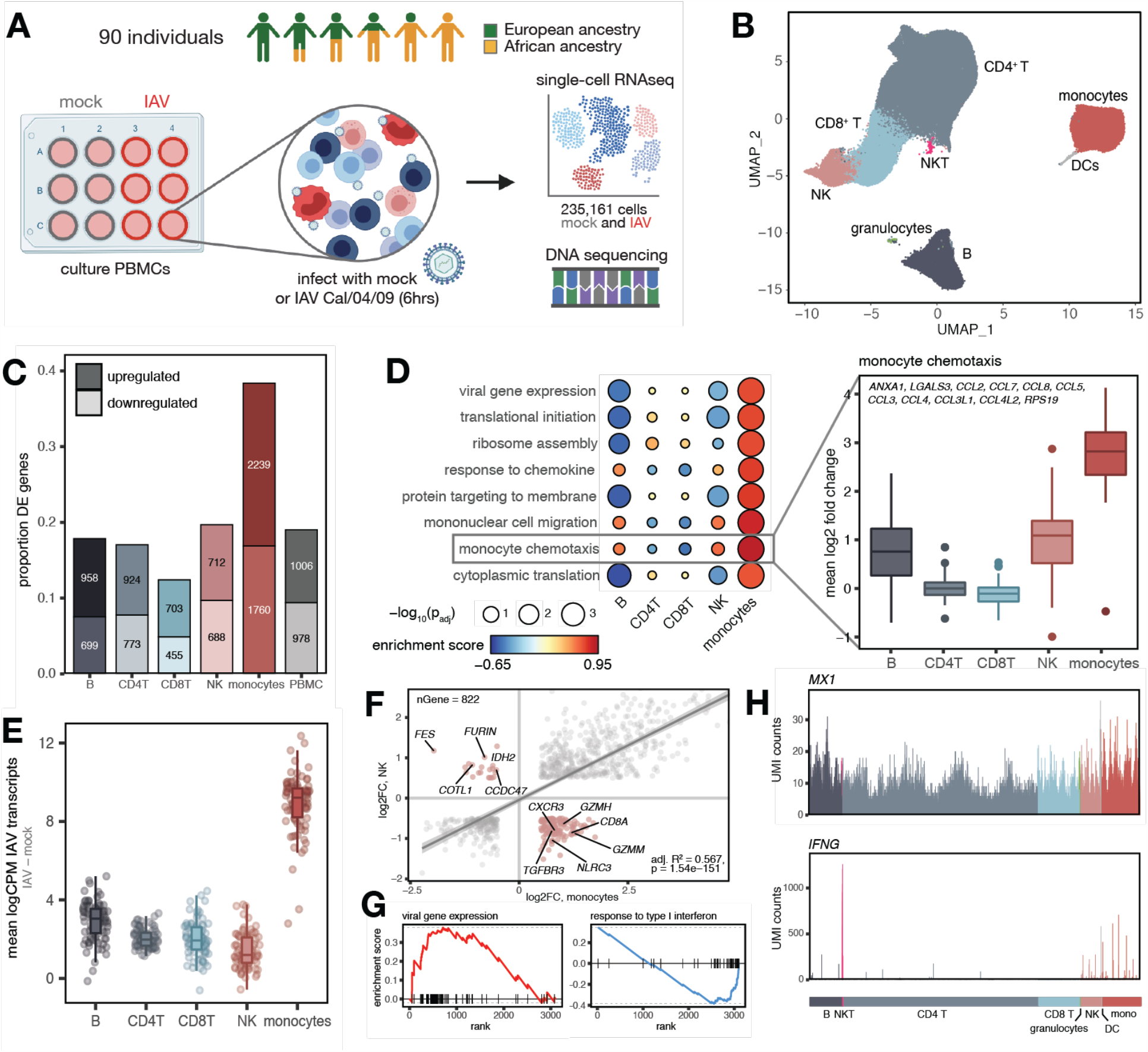
Shared and cell type-specific transcriptional responses to IAV infection. (A) Study design schematic. PBMCs from 90 individuals were exposed to mock-conditioned media or IAV Cal/04/09 *in vitro* for 6 hours, followed by single-cell RNA sequencing and DNA collection for DNA sequencing. (B) UMAP of 235,161 high-quality single-cell transcriptomes from both mock-and IAV-infected cells across all individuals. (C) Numbers and proportions of genes that show differential expression (logFC > 0.5, FDR < 0.05) between mock- and IAV-infected conditions across the five major PBMC cell types. (D) Monocyte-specific GO pathways that show significant upregulation (enrichment score > 0, FDR < 0.10) following infection. Genes in the “monocyte chemotaxis” term are significantly more upregulated after IAV infection in monocytes compared to the other cell types (plotted means for each individual across genes in the IAV condition minus the mock condition, t-test, all p-values < 1×10^−10^ when comparing monocytes against each other cell type). (E) Distribution of IAV transcript expression across cell types, with monocytes showing a 3 – 6-fold higher number of IAV transcripts compared to any other cell type (t-test, all p-values < 1×10^−10^ when comparing monocytes against each other cell type). (F) Correlation between IAV infection effect sizes (log2 fold-change values) in monocytes (x-axis) and NK cells (y-axis) among DE genes in both monocytes and NK cells (n = 822). Line shows the best-fit slope and intercept from a linear model. Highlighted genes (pink) display discordant responses following IAV infection. (G) Example pathways enriched among genes with high (right, viral gene expression FDR = < 1×10^−10^) and low (left, response to type I interferon FDR = 7.1×10^−4^) specificity scores, where all genes are rank-ordered by specificity score on the x-axis (highest to lowest). (H) UMI counts (y-axis) per cell (x-axis) in the IAV-infected condition for example genes that show ubiquitous expression across cell types (*MX1*, top) and highly cell type-specific expression patterns (*IFNG*, bottom).

We first investigated the overall signature of IAV infection on our samples by collapsing the single-cell gene expression values for each of the five main clusters and for all cells together (i.e. “PBMCs”, including the major and minor clusters) into pseudobulk estimates per sample. Principal component analysis (PCA) on the PBMC pseudobulk expression data revealed a strong signature of the IAV infection effect, such that mock- and IAV-infected samples strongly separate on PC1, which explains 43% of the variance in the dataset (Fig. S1B, paired t-test, p < 1×10^−10^). Across the five main clusters and the total PBMC pool, monocytes were by far the most responsive to IAV infection (n = 3,999 differentially expressed (DE) genes [38.4% of those tested] with log2 fold-change > 0.5, FDR < 0.05). All other cell types displayed weaker infection effects (12.4 – 19.7% DE genes) (Fig. 1C, Table S2). In support of a major role for monocytes in the IAV response, gene set enrichment analysis (GSEA) revealed that genes that were more responsive in monocytes compared to all other cell types were strongly enriched for genes involved in viral transcription and monocyte-associated biological pathways, such as monocyte chemotaxis (FDR = 4.6×10^−4^) (Fig. 1D, Table S3). Moreover, monocytes exhibited the highest levels of intracellular IAV transcripts (i.e., influenza-derived transcripts generated and processed by infected host cells; > 3-fold increase compared to all other cell types, t-test, all p-values < 1×10^−10^ for monocytes compared to all other cell types) (Fig. 1E). This observation shows that monocytes are either the cell type most susceptible to viral entry, the most permissible to intracellular replication of the virus, or both.

We then explored the extent to which the infection response was concordant or discordant across the five major PBMC cell types. Overall, we found a strong correlation between the response to IAV infection across cell types (Pearson’s r range 0.65 – 0.95 across pairwise cell type comparisons, Fig. S1C). However, we also observed many genes for which the response to infection was discordant between cell types. For example, among differentially-expressed genes shared by monocytes and NK cells (n = 822), 138 genes (16.8%, Fig. S1D) responded to IAV infection in the opposite direction (Fig. 1F). To further dissect cell type-specific versus shared responses, we generated a per-gene specificity score based on the coefficient of variation of the log2 fold-change response across cell types for each gene that was significantly differentially-expressed in at least one cell type (high values indicate highly cell type-specific responses to IAV, low values indicate shared responses to IAV) (Table S4). Genes with highly cell type-specific patterns of response were enriched for roles in translational initiation, co-translational protein targeting to membrane, and viral gene expression (FDR < 1×10^−10^ for all terms, Fig. 1G, left, Table S4). In contrast, genes with low specificity scores were enriched for pathways related to type I interferon (IFN) signaling (FDR < 1×10^−10^) and response to type I IFN (FDR = 7.1×10^−4^) (Fig. 1G, right, Table S4). Thus, induction of IFN-related genes appears to be a fundamental component of the antiviral response that is shared across immune cell types (Fig. 1H, top). One notable exception to this observation is the gene *IFNG*, which encodes the type II IFN cytokine IFN-γ. IFN-γ is a crucial mediator of antiviral immunity (Kang et al., 2018), but shows an expression pattern that is almost exclusive to NKT cells in the IAV-infected condition (Fig. 1H, bottom). Collectively, our results underscore the importance of considering the immune responses of single cell types independently. Not only does this approach reveal distinct, cell type-specific responses to viral infection, but also highlights responses that would be undetectable or potentially misleading in more heterogeneous immune cell populations (e.g. PBMCs).

### Genetic ancestry is associated with the transcriptional immune response to IAV

We next identified genes for which gene expression levels are correlated with quantitative estimates of genetic ancestry in either the mock condition, the IAV-infected condition, or both (after controlling for age, batch, and other technical covariates). To increase power and improve our effect size estimates for these “population differentially-expressed” (popDE) genes, we implemented a multivariate adaptive shrinkage method (mash) (Urbut et al., 2019), which leverages the correlation structure of genetic ancestry effect sizes across cell types (see Methods for details in the statistical models used). Across conditions and cell types, we identified 1,949 popDE genes (local false sign rate (lfsr) < 0.10), ranging from 830 in NK cells to 1,235 genes in CD4^+^ T cells (Fig. 2A, Table S5). Within each cell type, most popDE genes were shared between mock and IAV-infected conditions (52.9% in monocytes – 77.4% in CD8^+^ T cells). In contrast, across cell types, we found that genetic ancestry effects on gene expression were highly cell type-specific, such that the majority of popDE genes were identified in only one or two cell types (52.2% in mock, 51.4% in IAV-infected) (Fig. 2B). For example, *CXCL8*, which encodes IL-8, an important mediator of the inflammatory response (Bickel, 1993), is more highly expressed with increasing African ancestry following IAV infection only in monocytes (lfsr = 0.051 in monocytes and lfsr > 0.25 in all other cell types) (Fig. 2C, top). There are, however, a minority of genes that exhibit shared genetic ancestry effects across all five cell types (17.8% in mock, 24.7% in IAV-infected). One such gene is *IL32*, which encodes a cytokine that induces other proinflammatory cytokines to activate the NF-κB pathway (Ca and Sh, 2006; Yan et al., 2018). *IL32* is more highly expressed with increasing African ancestry in all cell types following infection (lsfr < 8.8×10^−6^) (Fig. 2C, bottom).

**Fig. 2.**
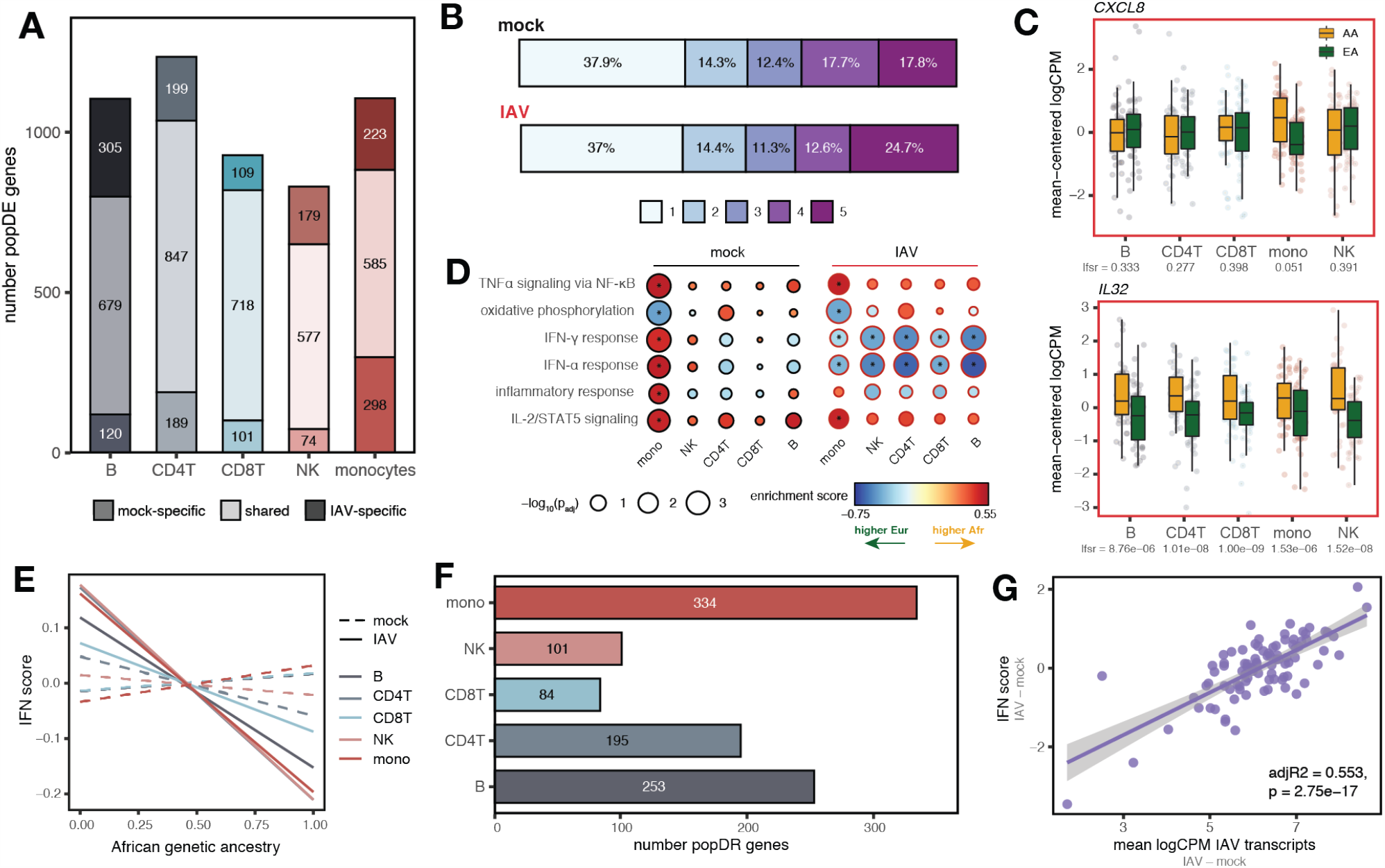
Genetic ancestry influences the immune response to IAV infection. (A) Number of significant mock-specific, shared, and IAV-specific popDE genes (lfsr < 0.10) across cell types. (B) Cell type sharing of significant popDE effects (1 = popDE effect is detected in only a single cell type, 5 = popDE effect is detected across all cell types). (C) Examples of cell type-specific (*CXCL8*, monocytes lfsr = 0.051, lfsr > 0.25 in all other cell types) and shared (*IL32*, lsfr < 8.8×10^−6^ in all cell types) popDE genes (AA in yellow, EA in green) in the IAV-infected condition. (D) GO enrichments for popDE effects across cell types in the mock-infected (black circles) and the IAV-infected (red circles) conditions. A positive enrichment score (ES) corresponds to an enrichment in genes with higher expression in individuals with increased African ancestry, while a negative ES corresponds to an enrichment in genes with higher expression in individuals with increased European ancestry. * represents pathways with FDR < 0.10. (E) Correlation between the proportion of African genetic ancestry (x-axis) and IFN score (y-axis) in the mock-infected (dotted lines, mean Pearson’s r across cell types = −0.0045, Fisher’s meta-p = 0.746) and the IAV-infected condition (solid lines, mean Pearson’s r across cell types = −0.26, Fisher’s meta-p = 2.9×10^−6^). (F) Number of significant popDR genes (lfsr < 0.10) across cell types. (G) IAV transcript levels (x-axis) predict the IFN score response (difference in IFN score between the IAV-infected and mock-infected conditions, y-axis) in PBMCs (adj R^2^ = 0.553, p = 2.8×10^−17^), and most individual cell types (Fig. S2C). In (E) and (G), lines show the best-fit slope and intercept from a linear model.

To identify the functional pathways most closely associated with genetic ancestry, we performed enrichment analysis on the popDE effects using the Molecular Signatures Database hallmark gene sets (Liberzon et al., 2015) (Fig. 2D, Table S6). In monocytes, we identified significant enrichment for multiple immune pathways prior to infection, including the IFN-α response (FDR = 1.9×10^−3^), IFN-γ response (FDR = 5.4×10^−4^), TNFα signaling via NF-κB (FDR = 6.1×10^−4^), IL-2/STAT5 signaling (FDR = 2.1×10^−3^), and inflammatory response (FDR = 0.012) (Fig. 2D). In all cases, these enrichments were identified for genes that were more highly expressed at baseline (i.e., in the mock treatment condition) in individuals with a greater proportion of African ancestry. Intriguingly, in IAV-infected cells, this pattern shifts: post-infection, we observed enrichment of type I and II IFN pathways (IFN-α response FDR = 0.014, IFN-γ response FDR = 0.040 in monocytes) (Fig. 2D) in genes more highly expressed with increasing European ancestry, as opposed to African ancestry. To better characterize this shift, we constructed a per-individual, per-condition score of interferon signaling activity (termed IFN score) by calculating the average mean-centered expression of genes belonging to the hallmark IFN-α and IFN-γ gene sets (Liberzon et al., 2015). This simple summary statistic revealed that increased European ancestry strongly correlates with increased IFN score, but only in the IAV condition (mean Pearson’s r across cell types = −0.26, Fisher’s meta-p = 2.9×10^−6^ in the IAV condition, Fisher’s meta-p = 0.746 in the mock condition) (Fig. 2E).

These findings suggest that, for some immune pathways (particularly interferon signaling pathways), genetic ancestry may also predict the magnitude of the response to IAV infection. To explicitly test this possibility, we identified significant interactions between treatment condition (mock versus IAV) and genetic ancestry levels. After mash estimation of interaction effect sizes across cell types, we identified 609 genes for which ancestry was associated with the response to infection (i.e., “population differentially-responsive” [popDR] genes, lfsr < 0.10). PopDR genes were found for all five cell types (number of popDR genes range = 84 – 334), but were most common in monocytes (n popDR genes = 334) (Fig. 2F); a core set of 27 popDR genes were also shared across cell types (Fig. S2A, Table S7). In agreement with our previous results, we found that increased European genetic ancestry predicts a stronger type I/II IFN response (measured as the difference in IFN score between the IAV-infected and mock conditions per individual) across cell types (mean Pearson’s r across cell types = −0.23, Fisher’s meta-p = 6.×10^−5^) (Fig. S2B). This observation cannot be explained by differences in baseline levels of IAV-specific serum IgG antibodies (Figure S2C, S2D). Type I/II IFN response magnitude also correlated with the level of IAV transcripts measured in PBMCs, such that the stronger the IFN response, the higher the measured IAV transcripts (adj. R^2^ = 0.553, p = 2.8×10^−17^) (Fig. 2G), a relationship primarily driven by monocytes (Fig. S2E). Accordingly, in line with their stronger type I/II IFN response, individuals with increased European ancestry proportion displayed increased levels of IAV transcript expression compared to individuals with higher levels of African ancestry (Pearson’s r = −0.32, p = 0.002, Fig. S2F), an observation that is replicated when monocytes are infected with IAV in isolation (O’Neill et al., 2020). These results point to the possibility that ancestry-associated variation in susceptibility to intracellular infection and/or differences in the ability to restrict viral replication may explain ancestry-associated differences in the type I/II IFN response.

### *Cis*-regulatory genetic variation explains ancestry-associated differences in gene regulation

To assess the contribution of genetic variation to genetic ancestry-associated differences in the transcriptional response to IAV infection, we integrated genome-wide expression profiles with genotyping data to map expression quantitative trait loci (eQTL) in both the mock and IAV-infected samples. We focused specifically on *cis*-eQTL, which we defined as SNPs located either within or flanking (±100 kilobases, kb) each gene of interest. We identified 2,234 genes that were associated with at least one *cis*-eQTL (lfsr < 0.10, hereafter referred to as eGenes) across all cell types and conditions tested (Fig. 3A, Table S8). Although many variants exert similar effects across cell types and infection conditions (45%, Fig. S3A), 13 - 24% of the eGenes identified within each cell type were only detected in one condition even after probing patterns of shared effects using mash (Urbut et al., 2019). Our results thus highlight the importance of gene– environment interactions in influencing transcriptional regulation in the immune system (Barreiro et al., 2012; Fairfax et al., 2014; Lee et al., 2014; Nédélec et al., 2016; Quach et al., 2016). Of note, we identified a set of 29 eGenes (Fig. S3A) that, across all cell types, were only detected in the IAV-infected condition, including the key IFN-inducible genes *OAS1, IFI44L, IFIT1, IRF1*, and *ISG15*. In *OAS1*, the top *cis* SNP across cell types (rs10774671) is an IAV-specific response eQTL (Fig. 3B, top: mock-infected lfsr = 0.81, bottom: IAV-infected lfsr = 1.5×10^−12^ in CD4^+^ T cells) that lies within a Neanderthal-derived haplotype (Sams et al., 2016) and that has been associated with higher OAS1 enzymatic activity (Bonnevie-Nielsen et al., 2005) and susceptibility to different viruses from the *Flaviviridae* family (El Awady et al., 2011; Kwon et al., 2013; Lim et al., 2009).

**Fig. 3.**
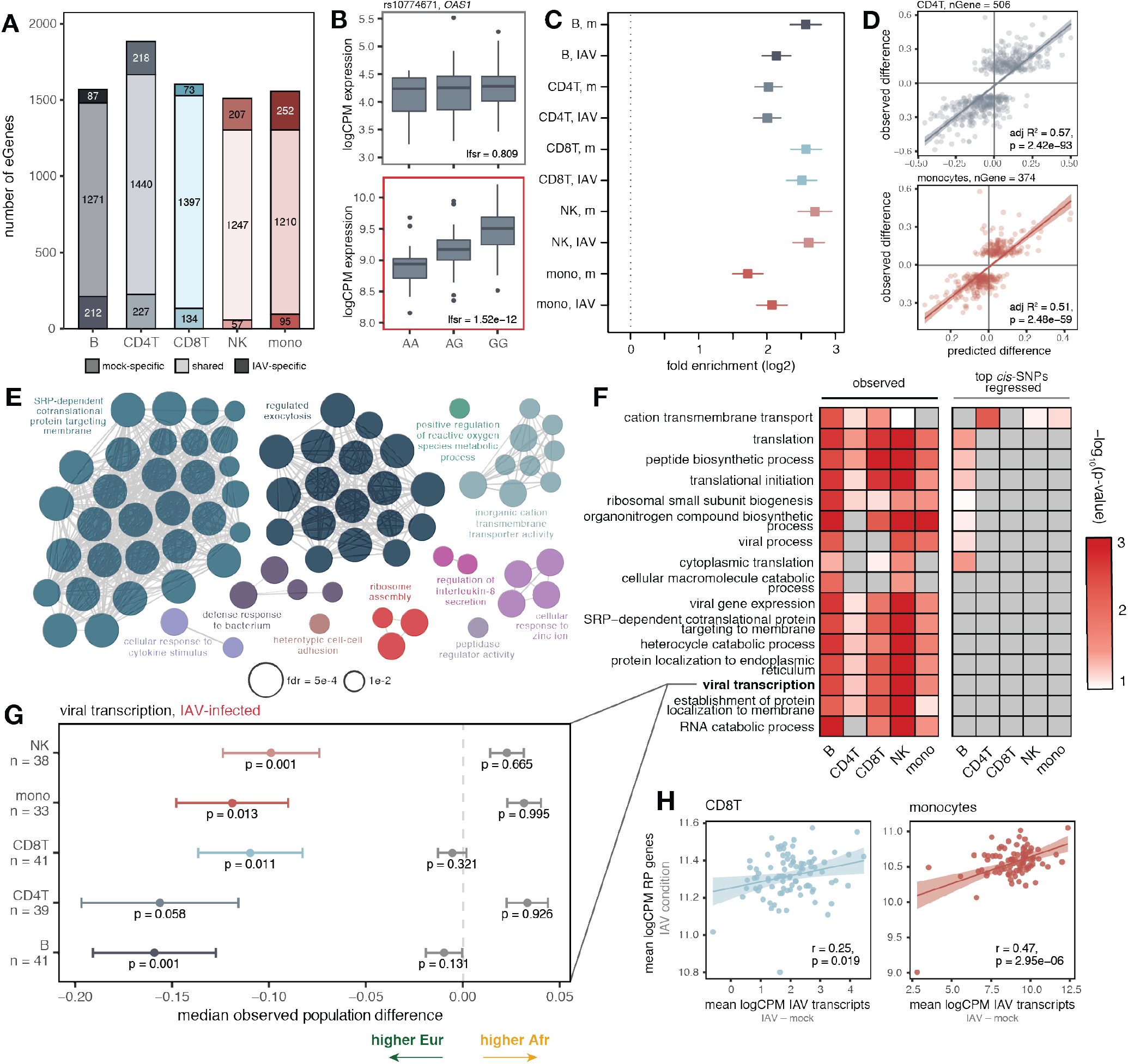
*Cis*-regulatory variation drives differences in the antiviral response, both at the individual and population levels. (A) Number of significant mock-specific, shared, and IAV-specific eGenes (lfsr < 0.10) across cell types. (B) Example of a condition-specific response eQTL (rs10774671 in *OAS1*) in CD4^+^ T cells (top: mock-infected, lfsr = 0.809, bottom: IAV-infected, lfsr = 1.5×10^−12^). (C) Enrichment of significant eGenes (lfsr < 0.10) among significant popDE genes (lfsr < 0.10) identified in each cell type and condition (x-axis: log2 fold enrichment with a 95% confidence interval; “m” = mock). (D) Correlation of the *cis*-predicted population differences in expression (x-axis) versus the observed population differences in expression (y-axis) among popDE genes with an eQTL in CD4^+^ T cells (top, adj R^2^ = 0.57, p = 2.42×10^−93^) and monocytes (bottom, adj R^2^ = 0.51, p = 2.48×10^−59^). (E) Significant (FDR < 0.01) ClueGO enrichments for the popDE genes that are also eGenes across all cell types in the IAV-infected condition. (F) Median observed population differences among genes in (E) for selected terms using a model estimating the observed genetic ancestry effects (left) and a model estimating this effect with the effect of the top *cis*-SNP regressed for all genes contained in the term (right). (G) Example term showing the effect of *cis* SNP regression. European-ancestry individuals display higher expression (median observed pop. difference < 0, colored point +/- SE) for the genes belonging to the “viral transcription” term in the observed data. Following *cis*-SNP regression (grey point +/- SE), this difference is attenuated. (H) Correlation between IAV transcript levels and the mean logCPM among ribosomal protein (RP) eGenes per individual in the IAV condition minus the mock condition for CD8^+^T cells (Pearson’s r = 0.25, p = 0.019) and monocytes (Pearson’s r = 0.47, p = 3×10^−6^). In (D) and (H), line shows the best-fit slope and intercept from a linear model.

We next tested whether eGenes were also likely to be differentially-expressed by genetic ancestry. Across all cell types and conditions, eGenes (lfsr < 0.10) were 3.2 to 6.5-fold more likely to be classified as popDE (lfsr < 0.10) than expected by chance (Fig. 3C). This enrichment suggests that ancestry-associated differences in gene expression are likely to have a substantial genetic component, perhaps due to divergence in allele frequencies at the causal eQTL. To formally evaluate this contribution, we calculated the correlation between 1) the estimated genetic ancestry effect from our popDE analysis, and 2) the predicted genetic ancestry effect based only on the effect size of the top eQTL per eGene and the genotype for this SNP among individuals of European and African ancestry (restricted to those popDE genes that were also eGenes in at least one cell type, n = 835 genes; see Methods for details). We found a striking correlation (Fig. 3D), such that genotype information on the eQTL alone explained an average of 52.5% (mock) and 53.6% (IAV-infected) of the variance in genetic ancestry effect sizes across cell types (Fig. S3B). These results indicate that, among popDE genes with an eQTL, on average, over 50% of the population differences are explained by *cis*-regulatory variation.

### Polygenic selection has shaped genetic ancestry-associated differences in ribosomal protein gene expression

We next sought to evaluate if the intersection set of popDE genes and eGenes clustered into specific biological pathways. We identified a strong enrichment for Gene Ontology (GO) terms related to transcriptional and translational processes, including “ribosomal small subunit biogenesis” and “viral transcription” (FDR < 3×10^−10^ in mock-infected and IAV-infected) (Fig. 3E), as well as more canonical immune-related pathways, such as myeloid/leukocyte activation and degranulation (Table S9). The observed gene expression divergence between populations in genes linked to similar biological functions could be explained by two hypotheses: 1) genes associated with such biological processes have evolved under relaxed evolutionary constraint, allowing them to accumulate *cis*-regulatory variants that have randomly diverged in allele frequencies via neutral genetic drift, or 2) *cis*-variants regulating these genes have undergone non-neutral shifts in allele frequencies, resulting in the accumulation of alleles that systematically influence the behavior of enriched pathways in a directional manner – a pattern consistent with polygenic selection.

To test for polygenic selection, for each of the enriched pathways, we calculated the median ancestry-associated differential expression effect (i.e., the estimated difference in gene expression between European- and African-ancestry individuals) across all popDE genes contained in a given pathway (limited to those with an eQTL). Under neutrality, we expect the direction of the ancestry-associated effects to be randomly distributed (i.e., some genes will be more highly expressed in European-ancestry individuals whereas others will be more highly expressed in African-ancestry individuals). In contrast, under polygenic selection, we expect to find a directional effect, such that most genes for a given pathway show higher expression in one ancestry group versus the other. Interestingly, most of the GO terms for ribosomal protein-related pathways (e.g. ribosomal biogenesis, viral transcription, etc.) show median population-associated differences in gene expression levels that are consistently higher in individuals with increased European ancestry across cell types, in both IAV-infected cells (Fig. 3F, 3G) and mock-exposed cells (Fig. S4A). Importantly, these differences are attenuated when regressing out the effects of the associated top *cis*-eQTLs for the genes (Fig. 3F, 3G), suggesting that such differences are driven by the cumulative effect of regulatory variants impacting the expression of ribosomal protein (RP) genes. Strikingly, we found a strong correlation between the average expression of RP eGenes and IAV transcript expression in both CD8^+^ T cells (Pearson’s r = 0.32, p = 0.002) and monocytes (Pearson’s r = 0.58, p < 1×10^−10^, Fig. 3H). Together, these data raise the possibility that viral infection-induced selection pressures have shaped ribosomal biology phenotypes in human populations, with potential implications for viral control mechanisms.

### Natural selection and susceptibility to autoimmune disease

Past selection imposed by pathogen exposure has been speculated to contribute to present-day susceptibility to autoimmune and chronic inflammatory diseases (Brinkworth and Barreiro, 2014; Sanz et al., 2018). However, it remains unclear whether natural selection has also contributed to genetic ancestry-associated differences in vulnerability to these diseases. To address this question, we performed colocalization analysis between the union set of eQTLs detected across all cell types and conditions and 14 publicly available genome-wide association study (GWAS) hits for 11 autoimmune diseases (Table S10). Colocalized eQTLs are expected to be strongly enriched for causal drivers of variation in disease susceptibility across individuals. Across all autoimmune diseases, we colocalized eQTL in our study with a total of 95 GWAS variants (Fig. 4A, Table S10).

**Fig. 4.**
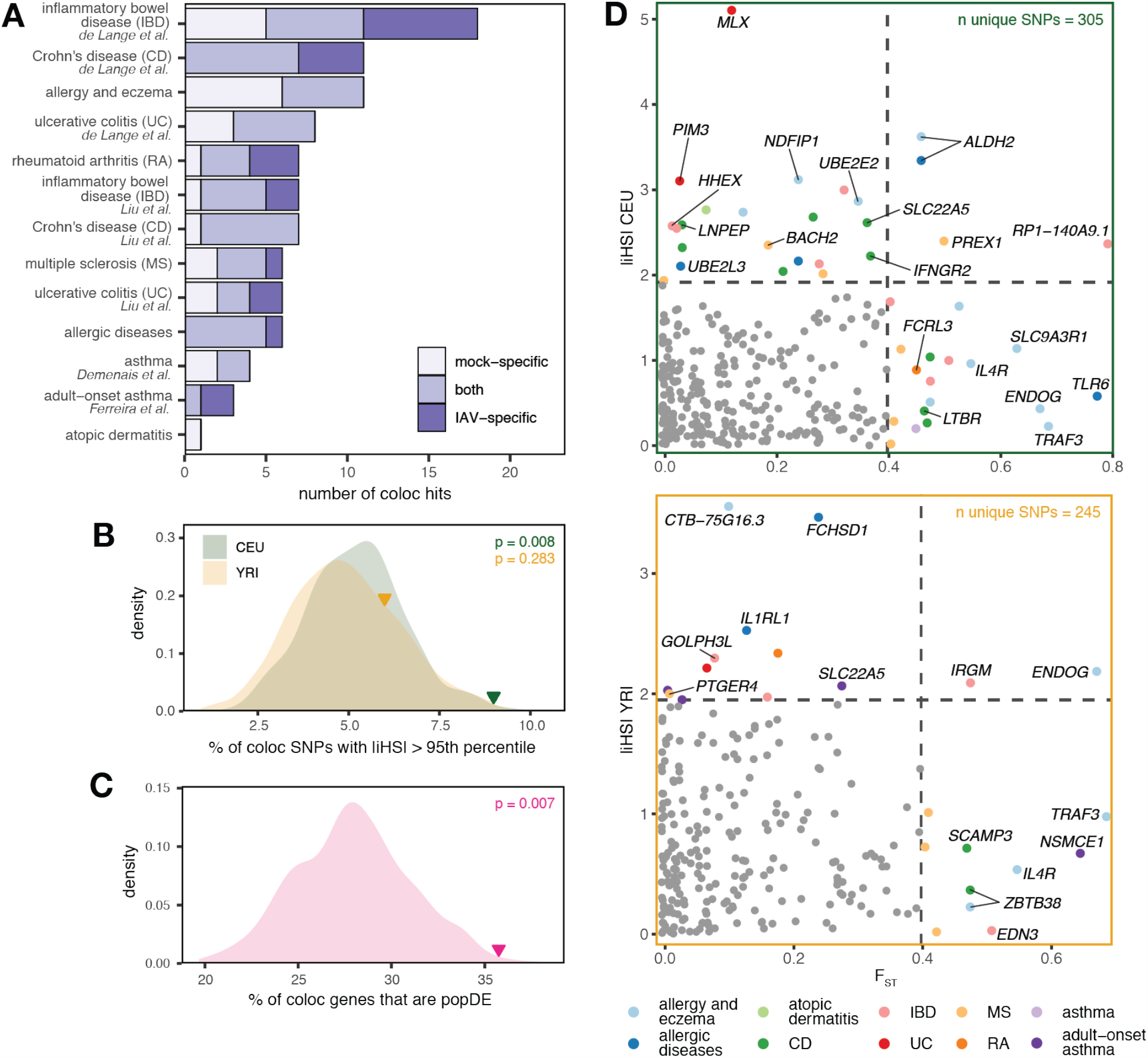
Recent positive selection has acted on *cis*-regulatory variants implicated in autoimmune disease risk. (A) Number of shared and condition-specific colocalization hits identified across cell types (x-axis) in the 11 autoimmune traits tested (y-axis). (B) Proportion of independent, colocalized lead GWAS loci that have |iHS| values > 95^th^ percentile of the genome-wide distribution among SNPs with > 5% MAF (CEU: green triangle, p = 0.008, YRI: yellow triangle, p = 0.283) compared to random expectation when sampling the same number of SNPs 1,000 times from all variants with a MAF > 5% in an LD-matched and MAF-matched manner (density distributions) among all autoimmune traits. (C) Proportion of genes with a colocalization signal that are popDE (pink triangle, p = 0.007) compared to random expectation when sampling the same total number of genes 1,000 times from all genes tested (density distribution) among all autoimmune traits. (D) F_ST_ and |iHS| values among the colocalized hits shown in A as well as those identified in the harmonized bulk eQTL data. F_ST_ values are plotted on the x-axis, while |iHS| values are plotted on the y-axis (top: CEU, bottom: YRI). Dotted lines show the 95^th^ percentile of the genome-wide distribution for the respective selection statistic (F_ST_ = 0.398, |iHS| CEU = 1.92, |iHS| YRI = 1.95). eGenes with a selection statistic > 95^th^ percentile are represented by a colored point, and colors represent the autoimmune trait for which a colocalization signal is detected (here, the multiple inflammatory bowel disease, ulcerative colitis, and Crohn’s disease GWAS have been collapsed into a single label).

To analyze a broader array of immune-related colocalization signals, we combined our data with colocalization results for bulk eQTLs in 18 immune cell types from 3 large immune eQTL studies (DICE (n = 91) (Schmiedel et al., 2018), DGN (n = 922) (Battle et al., 2014), and BLUEPRINT (n = 197) (Chen et al., 2016)) for the same 14 autoimmune GWAS (Mu et al., 2020). This approach allowed us to identify 1,030 colocalized GWAS hits across the 11 traits (mapping to 536 eGenes, Table S10). We then asked if these putative causal variants were enriched for signatures of natural selection in the 1000 Genomes Project CEU and YRI populations, using either the integrated haplotype score (iHS, a within-population measure of recent positive selection based on haplotype homozygosity (Voight et al., 2006)) or extreme values of population differentiation (F_ST_). Far more colocalized loci display high |iHS| scores (values > 95^th^ percentile of the genome-wide distribution) in the CEU population than expected by chance (p = 0.008, Fig. 4B), while no significant enrichment was detected in YRI. Our results thus suggest that natural selection has acted on these *cis*-regulatory autoimmune risk variants, particularly in Europeans, with the caveat that the vast majority of GWAS studies to date have focused exclusively on individuals of European ancestry (Bustamante et al., 2011), preventing us from detecting signatures of selection among GWAS loci unique to African-ancestry individuals. Moreover, we observed that colocalized genes are more likely to be differentially-expressed between populations than expected by chance (35.8% are classified as popDE, p = 0.007) (Fig. 4C), pointing to a potential genetic contribution for the differences in the incidence of autoimmune and inflammatory disorders reported between African and European-ancestry individuals (Brinkworth and Barreiro, 2014).

Within our set of colocalized eGene-SNP pairs, 48 eGenes carried a signature of recent positive selection in either the CEU or YRI populations (|iHS| or F_ST_ > 95^th^ percentile of the genome-wide distribution) (Fig. 4D). Many of these genes involve crucial immune-related functions. For example, the Crohn’s disease-susceptibility risk variant rs2284553 colocalized with *IFNGR2*, the gene encoding the beta chain of the IFN-γ receptor, in naïve CD8^+^ T cells (Fig. S5A). This variant is found at much higher frequency in the CEU population (MAF = 0.38) than the YRI population (MAF = 0.05) and shows a signature of recent positive selection in the CEU (iHS = 2.22). Another variant detected in the allergic disease GWAS, rs5743618, maps to a non-synonymous SNP located in *TLR1* that is also an eQTL for the nearby gene *TLR6* (Fig. S5B). This variant compromises NF-κB signaling and activation to produce an attenuated inflammatory response (Barreiro et al., 2009) and is a known *trans*-regulatory hotspot (Piasecka et al., 2018; Quach et al., 2016). Notably, it is found at low frequency in the YRI population (derived allele frequency (DAF) = 0.04) but is found at elevated frequency in the CEU population (DAF = 0.67) (Sanz et al., 2018). This difference in allele frequency alone explains the positive correlation between African genetic ancestry and the transcriptional response to immune stimulation with antigens that signal through TLR1 (Nédélec et al., 2016; Quach et al., 2016; Sanz et al., 2018).

## Discussion

Together, our results provide the most detailed characterization to date of the genetic determinants that shape inter-individual and genetic ancestry-associated differences in the response to viral infection across the five most common immune cell types found in PBMCs. We identified thousands of genes for which expression levels are correlated with genetic ancestry across different immune cell types, but found that the majority of these cases (52.2% in mock, 51.4% in IAV-infected) are restricted to only one or two cell types. These results are likely explained by a combination of cell type-specific genetic effects and environmental factors that correlate with genetic ancestry but that only act on certain cell types. For example, chronic stress has been shown to causally alter immune gene regulation, yet its effects are mainly limited to helper T cells and NK cells (Snyder-Mackler et al., 2016). Although our findings corroborate previous reports of elevated inflammatory pathway activity with increasing African ancestry, at least at baseline (*25, 26*), they also reveal a novel pattern: increased activity of type I/II IFN pathways following influenza infection associated with increased European ancestry. This observation has potential clinical implications, as interferons are the main defensive cytokines released during the early phase of acute influenza infection as well as most other viral infections. However, when chronically elevated, interferons can increase susceptibility to the uncontrolled inflammation typical of severe cases of influenza and now COVID-19 (Vabret et al., 2020). More studies are now needed to define whether ancestry-associated variation in the interferon response to viral infection *in vivo* is associated with differential viral clearance, disease severity, and disease outcome.

Many of the genetic ancestry-associated differences in immune regulation we observe are driven by allele frequency differences at *cis*-regulatory variants. Among popDE genes in which we identify at least one *cis*-eQTL across cell types and conditions, we estimate that, on average, *cis*-eQTLs explain approximately 53% of the variance in the observed ancestry-associated differences. This is likely an underestimate, given that it assumes that each gene has only a single *cis*-eQTL, when in fact many genes have been shown to have two or more independent cis-eQTL (Lappalainen et al., 2013). Further, we are underpowered to detect *trans* associations, and we do not consider non-SNP regulatory variants (e.g., indels and copy number variation), which also influence gene expression variation in humans (Gymrek et al., 2016). In addition, we provide evidence for ancestry-associated directional shifts in molecular traits (i.e., gene expression phenotypes related to specific biological pathways) that are under *cis*-regulatory genetic control, highlighting the potential role of polygenic selection in the history of these phenotypes.

The signature of selection at ribosomal protein (RP) genes (*RPL, RPS*) is of particular interest, as RPs facilitate translation initiation of viral transcripts (Haque and Mir, 2010; Huang et al., 2012) and directly interact with viral mRNA and proteins to enable viral protein synthesis (Li, 2019). These proteins also play essential roles in ribosomal biogenesis (Fromont-Racine et al., 2003), a process that influences viral reproduction and cell-intrinsic immune responses following human cytomegalovirus infection (Bianco and Mohr, 2019), and that also affects innate immune signaling pathways to modulate the IFN-γ-mediated inflammatory response (Vyas et al., 2009) and NF-κB target gene expression (Wan et al., 2011). Further, a subset of ribosomes, known as immunoribosomes, has been hypothesized to preferentially synthesize antigenically-relevant cellular and viral peptides for immunosurveillance by the MHC class I system, resulting in a tight link between translation and antigen presentation that may allow immune cells to more quickly recognize and eliminate infected cells (Wei and Yewdell, 2019; Yewdell, 2007). Together, these observations support the argument that RPs are important mediators of the host immune response to virus, and raise the possibility that polygenic selection on ribosomal pathways has contributed to present-day variation in viral control within and between human populations.

Finally, our results provide evidence that recent, local positive selection has acted on putatively causal regulatory risk variants associated with common autoimmune diseases in GWAS, strengthening the link between pathogen-mediated selection and susceptibility to autoimmune disorders (Brinkworth and Barreiro, 2014; Nielsen et al., 2017; Quach and Quintana-Murci, 2017). The connection between infectious diseases and chronic inflammatory disorders is further supported by reports that some pathogens are contributing, and possibly causal, factors to the development of certain chronic inflammatory and autoimmune diseases (e.g., Epstein–Barr virus and systemic lupus erythematosus, rheumatoid arthritis, and multiple sclerosis; *Mycobacterium avium* and Crohn’s disease; *Yersinia enterocolica* and inflammatory bowel disease) (Abubakar et al., 2008; Feller et al., 2007; James et al., 2001; Ramos-Casals et al., 2005; Saebo et al., 2005; Yamazaki et al., 2005). Our findings shed light on human evolutionary history and lend key empirical support to arguments that link historical pathogen-mediated selection to present-day susceptibility to autoimmune and inflammatory diseases.

## Supporting information

Table S1: Sample meta data

Table S2: Global infection effects

Table S3: Global infection DE enrichments

Table S4: Ranked specificity scores and enrichments

Table S5: PopDE effects

Table S6: PopDE effect enrichments

Table S7: PopDR effects

Table S8: eQTL effects

Table S9: GO enrichments for popDE genes with an eQTL

Table S10: GWAS autoimmune traits and colocalization results

## Acknowledgements

We thank Jenny Tung, Briana Mittleman, Genelle Harrison, and members of the Barreiro lab for their constructive comments and feedback. This work was completed in part with resources provided by the University of Chicago Research Computing Center. Figure 1A was created with BioRender.com.

## Author contributions

L.B.B directed the study. H.E.R. and L.B.B designed the experiments. J.K.F, B.K.T., and R.A.L. generated the influenza A Cal/04/09 strain used for the infection experiments. H.E.R. performed the *in vitro* PBMC infection experiments. H.E.R. performed the sequencing library preparation, with help from M.S.C. H.E.R. led the computational analyses, with contributions from Z.M., Y.I.L. (colocalization analysis), J.C.G. and J.G.H. (iHS calculations). H.E.R. and L.B.B. wrote the manuscript, with input from all authors.

## Methods

### Peripheral blood mononuclear cell (PBMC) collections

All samples were obtained from BioIVT. A signed, written consent was obtained from each participant. Blood was collected from 90 male donors between the ages of 21 – 69 who identified as either African-American (AA) (n = 45) or European-American (EA) (n = 45) from the same collection site in Miami, Florida (United States) utilizing a standard protocol with a sodium heparin anticoagulant. Briefly, PBMCs were extracted from whole blood using a density gradient, washed with HBSS, reconstituted in CryoStor CS10 to a concentration of 10 million (M) cells/ml, and subsequently cryopreserved. Between 6 – 10M cells per individual were frozen per vial. We decided to only focus on males to avoid the potentially confounding effects of sex-specific transcriptional differences in the response to infection. Only individuals self-reported as currently healthy were included in the study. All individuals had detectable levels of IAV-specific serum IgG antibodies, but no differences in antibody titers were identified between European and African-ancestry individuals (Figure S2C, S2D).

### Generation of influenza A virus

Influenza A virus California/04/2009 (Cal/04/09) virus was rescued in 293T cells by plasmid-based transfection with IAV Cal/04/09 in the pDZ vector using methods previously described (Fodor et al., 1999; Hai et al., 2010; Hoffmann et al., 2000). 24 hours following transfection, 7.5×10^5^ MDCK cells were added to the culture in Opti-MEM containing TPCK trypsin (1 μg/mL). For the following two days, 500 μL of Opti-MEM containing TPCK trypsin (2 μg/mL) was added to the culture. One day later, the supernatant was harvested, centrifuged to remove cellular debris, and stored at -80°C. Cal/04/09 was amplified on MDCK cells to generate a stock. Uninfected MDCK cells were cultured for 48 – 72 hours and supernatant was harvested to generate the control, mock-conditioned media. Stocks were plaqued on MDCK cells. Cells were infected in infection media (PBS with 10% Ca/Mg, 1% penicillin/streptomycin, 5% BSA) at 37°C for 1 hour. Infection media was replaced with an agar overlay (2X MEM, 1 μg/mL TPCK trypsin, 1% DEAE-dextran, 5% NaCo_3_, 2% oxoid agar), and cells were cultured at 37°C for 40 hours then fixed with 4% formaldehyde. Blocking and immunostaining were done for 1 hour at 25°C in 5% milk. Primary stain was mouse anti-Cal/04/09 (1:5000), secondary stain was peroxidase sheep anti-mouse-HRP (1:5000) (45001275, GE Healthcare). TrueBlue Peroxidase Substrate (50-647-28, Kirkegard & Perry Laboratories) was used as directed for detection of virus plaques.

### *In vitro* infection experiments and sample collections

PBMCs were unfrozen approximately 14 hours prior to infection and cultured overnight in RPMI 1640 supplemented with 10% fetal bovine serum (FBS), 2 mM L-glutamine, and 10 ug/ml gentamycin. Infection experiments were performed over 15 batches, where each experimental batch was semi-balanced for self-identified ancestry label to avoid introducing a batch effect confounded with genetic ancestry. The morning of the experiment, 1M PBMCs were plated at a concentration of 1M/ml for each condition, and exposed to either mock-conditioned media (negative control) or Cal/04/09 IAV at an MOI of 0.5. After 30 minutes of exposure, the control media or virus was washed from PBMC cultures, cells were replated, and cells were then incubated for 6 hours at 37°C in 5% CO_2_ and 20% O_2_. Following the 6 hour incubation, cells were collected, washed, and prepared for single-cell capture using the 10X workflow. Immediately prior to the capture, cells from samples were combined into two pools (6 samples per pool) each balanced for infection status (mock-infected and IAV-infected) and genetic ancestry (Table S1). Multiplexed cell pools were used as input for the single-cell captures, and for each cell pool, 10,000 cells were targeted for collection using the Chromium Single Cell 3’ Reagent (v2 chemistry) kit (10X Genomics). Post Gel Bead-in-Emulsion (GEM) generation, the reverse transcription (RT) reaction was performed in a thermal cycler as described (53°C for 45 min, 85°C for 5 min), and post-RT products were stored at -20°C until downstream processing (no longer than 4 days post-RT reaction). For DNA processing, 1M PBMCs were collected, and DNA was extracted using the DNeasy Blood and Tissue Kit (Qiagen) following the “Cultured cells” protocol.

### Single-cell library preparation and RNA-sequencing

Post-RT reaction cleanup, cDNA amplification, and sequencing library preparation were performed as described in the Single Cell 3’ Reagent Kits v2 User Guide (10X Genomics). Briefly, cDNA was cleaned with DynaBeads MyOne SILANE beads (ThermoFisher Scientific) and amplified in a thermal cycler using the following program: 98°C for 3 min, 11 cycles × 98°C for 15 s, 67°C for 20 s, 72°C for 1 min, and 72°C 1 min. After cleanup with the SPRIselect reagent kit (Beckman Coulter), the libraries were constructed by performing the following steps: fragmentation, end-repair, A-tailing, SPRIselect cleanup, adaptor ligation, SPRIselect cleanup, sample index PCR (98°C for 45 s, 14 cycles × 98°C for 20 s, 54°C for 30 s, 72°C for 20 s, and 72°C 1 min), and SPRIselect size selection. Batches of four experiments (corresponding to eight multiplexed single-cell captures) were processed at a time. Prior to sequencing, all multiplexed single-cell libraries (n = 30) were quantified using the KAPA Library Quantification Kit for Illumina Platforms (Roche) and pooled in an equimolar ratio. Libraries were sequenced 100 base pair paired-end (R1: 30 cycles, I1: 10 cycles, R2: 85 cycles) on an Illumina NovaSeq to an average depth of 45,612 mean reads per cell across all batches (average median genes detected per cell across batches = 689).

### Low-pass DNA sequencing and VCF processing

Out of the 90 individuals in the cohort, 89 were successfully genotyped using DNBseq low-pass whole-genome sequencing (BGI) at 4x coverage. Variants were called across individuals using the human reference genome (GRCh37), yielding a merged VCF, and the ImputeSeq low-pass imputation pipeline (Gencove) was used to perform VCF imputation. The imputed merged VCF was lifted over to GRCh38 with CrossMap (v0.3.9) (Zhao et al., 2014) using the GRCh37 to GRCh38 Ensembl chain file downloaded at ftp://ftp.ensembl.org/pub/assembly_mapping/homo_sapiens/ and the GRCh38 FASTA file from ftp://ftp.ensembl.org/pub/release-92/fasta/homo_sapiens/dna/. For each individual, low-quality variants were filtered by only retaining those with a maximum genotype probability (GP in FORMAT field) > 0.90 using QCTOOL (v2.0.7, https://www.well.ox.ac.uk/~gav/qctool_v2/). If the max(GP) for a variant was < 0.90, the variant call was automatically set to missing. Only autosomal, biallelic SNPs were kept for downstream analysis using the SelectVariants function (--select-type-to-include SNP) from GATK (v3.7).

### Estimation of genome-wide admixture levels

Prior to estimation of genome-wide admixture proportions, samples were merged with CEU (n = 99, Utah Residents [CEPH] with Northern and Western European Ancestry) and YRI (n = 108, Yoruba in Ibadan, Nigeria) samples from the 1000 Genomes Project (1000GP) Phase 3 dataset (Auton et al., 2015) (downloaded from ftp://ftp.1000genomes.ebi.ac.uk/vol1/ftp/release/20130502/supporting/GRCh38_positions/). The proportion of European and African genetic ancestry for each individual included in the study was estimated using the supervised clustering algorithm in ADMIXTURE (v1.3.0) (Alexander and Lange, 2011). A total of 13,518,147 unlinked SNPs (r^2^ between all pairs < 0.1) were used for genetic ancestry assignments, assuming k = 2 ancestral clusters. These estimated quantitative genetic ancestry proportions were used to assess differences in immune responses between populations.

### Mapping, demultiplexing, and initial cell filtering

FASTQ files from each multiplexed capture library were mapped to a custom reference containing GRCh38 and the Cal/04/09 IAV reference genome (downloaded from NCBI, created using cellranger mkref) using the cellranger (v3.0.2) (10X Genomics) count function. Souporcell (v2.0, Singularity v3.4.0) (Heaton et al., 2020) in --skip_remap mode (-k 6) was used to demultiplex cells into samples based on genotypes from a common variants file (1000GP samples filtered to SNPs with >= 2% allele frequency in the population, downloaded from https://github.com/wheaton5/souporcell). Briefly, souporcell clusters cells based on cell allele counts in common variants, assigning all cells with similar allele counts to a single cluster corresponding to one individual, while also estimating singlet/doublet/negative status for that cell. For each batch, hierarchical clustering of the true genotypes known for each individual (obtained from low-pass whole-genome-sequencing) and the cluster genotypes estimated from souporcell was used to assign individual IDs to souporcell cell clusters. All 89 individuals were successfully assigned to a single cluster.

After demultiplexing cells into samples, Seurat (v3.1.5, R v3.6.3) (Stuart et al., 2019) was used to perform quality control filtering of cells. In total, we captured 255,731 cells prior to filtering (range of cells recovered per capture: min. = 5534, max. = 10805). Cells were considered “high-quality” and retained for downstream analysis if they had: 1) a “singlet” status called by souporcell, 2) between 200 – 2500 genes detected (nFeature_RNA), and 3) a mitochondrial reads percentage < 10%, leaving 236,993 cells (n = 19,248 genes).

### Clustering, cell type assignment, and UMAP analysis

We performed two versions of clustering analysis and cell type assignment: 1) in which IAV genes were kept in the raw count matrix (used as input for pseudobulk calculations), and 2) in which IAV genes were subset out of the raw count matrix (for visualization of the UMAP in Fig. 1B). All other steps of the clustering workflow (implemented in Seurat v3.1.5) remained the same. Pseudobulk expression estimates (see below) between clustering versions for cell type-matched clusters were extremely similar (adj R^2^ > 0.999 for comparisons between versions). For both clustering iterations, we split the cells by infection status (mock or IAV) and ran SCTransform to normalize and scale the UMI counts within condition. In this step, we simultaneously regressed out variables corresponding to experiment batch and percent mitochondrial reads per cell. The data was then integrated on infection status using the SelectIntegrationFeatures, PrepSCTIntegration, FindIntegrationAnchors, and IntegrateData workflow. Following integration, dimensionality reduction was performed via UMAP (RunUMAP function, dims = 1:30) and PCA (RunPCA function, npcs = 30). A Shared Nearest Neighbor (SNN) Graph was constructed using the FindNeighbors function (dims = 1:20, all other parameters set to default), and clusters were subsequently called using the FindClusters algorithm (resolution = 0.5, all other parameters set to default).

Clusters were annotated based on the expression of canonical immune cell marker genes (CD4^+^ T: *CD3D*^*+*^, *CD3E*^*+*^, *CD8A*^*–*^; CD8^+^ T: *CD3D*^*+*^, *CD8A*^*+*^; NK cells: *CD3D*^*–*^, *NKG7*^*+*^, *GNLY*^*+*^; monocytes: *CD14*^+^, *LYZ*^+^; B: *MS4A1*^+^; granulocytes: *PRSS57*^+^; dendritic cells (DCs): *HLA-DRA*^+^, *HLA-DRB1*^+^, *CCR7*^+^, *CST3*^+^, *CD83*^+^). A small group of cells, which were identified as B cells, clustered with CD4^+^ T cells in the UMAP (Fig. 1B), and we investigated this further to see whether this subset represented a distinct, rare cell type. Further analysis revealed that these cells express markers typical of NKT cells, including *CD3D, NKG7, IL2, TNF*, and *IFNG*, and thus, these cells were manually annotated as NKT cells. In the UMAP constructed from input data containing IAV genes, we excluded 1,832 cells for which we could not confidently assign a cell type, as they clustered on the basis of high IAV transcript expression, leaving us with 235,161 cells across all individuals and conditions for downstream analysis (n CD4^+^ T cells = 138,801, CD8^+^ T cells = 32,446, monocytes = 27,020, B cells = 22,877, NK cells = 13,220, DCs = 374, granulocytes = 301, NKT cells = 122).

### Calculation of pseudobulk estimates

Cluster-specific pseudobulk estimates were used to summarize single-cell expression values into bulk-like expression estimates within samples (where, here, a sample is an individual, infection-condition pair, n = 180). This was performed for all five major cell types (CD4^+^ T cells, CD8^+^ T cells, B cells, monocytes, NK cells) and PBMCs, where all high-quality cells from all cell types identified (n = 235,161) were treated as a single aggregate cluster. Within each cluster for each sample, raw UMI counts were summed across all cells assigned to that sample for each gene using the sparse_Sums function in textTinyR (v1.1.3), yielding an n × m expression matrix, where n is the number of samples included in the study (n = 180) and m is the number of genes detected in the single-cell analysis (m = 19,248) for each of the 6 clusters.

### Calculation of capture-corrected expression for downstream modeling

From this point forward, pseudobulk estimates were treated as de facto bulk expression data for each cell type considered. As such, calculations of residuals and downstream modeling of infection and genetic ancestry effects (see below) were performed for each cluster independently. For each cell type, lowly-expressed genes were filtered using cell-type specific cutoffs (removed genes with a median logCPM < 1.5 in CD4^+^ T cells, monocytes, and PBMCs, < 2.5 in B cells and CD8^+^ T cells, and < 4.0 in NK cells), leaving the following number of genes per cell type: CD4^+^ T cells = 9,291, CD8^+^ T cells = 9,960, B cells = 9,335, monocytes = 10,424, NK cells = 7,109, and PBMCs = 10,430.

After removing lowly-expressed genes, normalization factors to scale the raw library sizes were calculated using calcNormFactors in edgeR (v 3.26.8) (Robinson et al., 2010). The voom function in limma (v3.40.6) (Ritchie et al., 2015) was used to apply these size factors, estimate the mean-variance relationship, and convert raw pseudocounts to logCPM values. A model evaluating the technical effect of capture (∼ 0 + capture, where capture corresponds to a factor variable representing the 30 experimental capture batches) on gene expression was fit using the lmFit and eBayes functions, and model residuals were obtained using the residuals.MArrayLM function in limma. The average capture effect was then computed by taking the mean of the capture coefficients across all 30 capture batches per gene, and this average capture effect was added back to the residuals across samples to generate the capture-corrected expression estimates. The inverse variance weights calculated by voom were obtained and included in the respective lmFit call for all downstream models unless otherwise noted.

While performing quality control checks on our data, we noticed that the density distributions of the capture-corrected expression estimates were bimodal for some samples in certain cell types. We estimated this bimodality proportion in each cell type for each sample by: i) estimating the local minimum of the density distribution, ii) subsetting the x-axis on a restricted range that was specific to each cell type, iii) using the × value where y equals the estimated local minimum as the bimodal threshold, and iv) calculating the proportion of genes less than this threshold. Assigned bimodality proportions were manually checked and corrected to an approximate value if they were obviously over- or under-estimated. The bimodality proportion is negatively correlated with cell counts per sample in most cell types and was most pronounced in the CD8^+^ T cells, monocytes, and NK cells (i.e. the cell types with the fewest number of cells collected per sample). To remove any potentially confounding effects associated with this artifact, the appropriate cell-type specific bimodality proportion vector across samples was included as a quantitative technical covariate in all of our downstream models.

### Modeling global infection effects

To obtain estimates of the global infection effects, capture-corrected expression levels of samples corresponding to the same individual were compared in a paired design, in which individuals were introduced as additional covariates into the following differential infection effect model that was run per cell type:

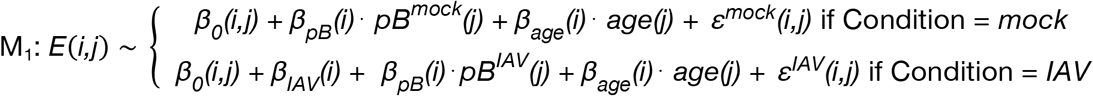

Here, *E (i,j)* represents the capture-corrected expression estimate of gene *i* for individual *j* and *β*_*0*_*(i,j)* represents the intercept corresponding to gene *i* and individual *j* (i.e. the expectation of gene *i*’s expression level in the mock-infected sample for individual *j*). When evaluated, this model gives the global estimate of the IAV infection effect per gene, *β*_*IAV*_*(i)*, approximated using the within-individual variation in gene expression across conditions. Further, *pB*^*cdt*^ represents the bimodal proportion estimated per sample for the respective cell type being modeled (where *cdt* represents either the mock or IAV), with *β*_*pB*_ being the corresponding effect on gene expression, and age represents the mean-centered, scaled (mean = 0, sd = 1) age in years per individual, with *β*_*age*_being the impact of age on expression. Finally, *εcdt* represents the residuals for each respective condition (mock or IAV) for each gene *i*, individual *j* pair. Of note, when modeling the expression estimates in PBMCs, two additional covariates were added to the model, corresponding to the first two principal components of a PCA performed on an n × m cell type proportion matrix (where n = number of samples = 180, m = number of cell types = 10, with the matrix populated by the cell type proportions for each sample [calculated by the number of cells per cell type cluster for a sample divided by the total number of cells assigned to that sample]) to account for the majority of the variance introduced by underlying cell type composition (PC1 percent variance explained (PVE) = 53.8%, PC2 PVE = 23.2%, total = 77.0%).

These models were fit using the lmFit and eBayes functions in limma (Ritchie et al., 2015), and the estimates of the global infection effect *β*_*IAV*_*(i)* (i.e. the differential expression effects due to IAV infection) were extracted across all genes along with their corresponding p-values. We controlled for false discovery rates (FDR) using an approach analogous to that of Storey and Tibshirani (Nédélec et al., 2016; Storey and Tibshirani, 2003), which makes no explicit assumptions regarding the distribution of the null model but instead derives it empirically. To obtain a null, we performed 10 permutations, where infection status label (mock/IAV) was permuted within individual. We consider genes significantly differentially-expressed upon infection if they have a *β*_*IAV*_ |logFC| > 0.5 and an FDR < 0.05.

### Calculation of IFN score

To construct the IFN score metric, we summarized the expression patterns of genes involved in the type I/II IFN response as a whole, where, within condition, we i) subset on genes belonging to the hallmark IFN gamma and alpha response pathways, ii) mean-centered and scaled the expression values for each gene across individuals, and iii) computed the average scaled expression across genes per individual.

### Modeling genetic ancestry effects and integration with mashr

Prior to modeling genetic ancestry effects, capture-corrected expression estimates were quantile-normalized within condition using qqnorm in R. The following nested linear model was used to identify genes for which expression levels are correlated with the proportion of African ancestry across individuals within condition (i.e. popDE genes):

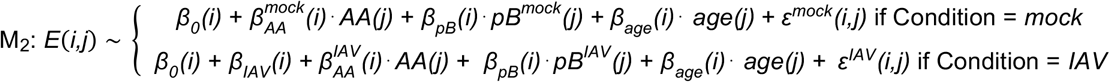

Here, *E (i,j)* represents the capture-corrected expression estimate of gene *i* for individual *j, β*_*0*_*(i)* is the global intercept accounting for the expected expression of gene *i* in a 100% European-ancestry mock-infected individual, 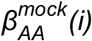 and 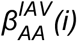 indicate the effects of African admixture (mean-centered, scaled African ancestry proportion, *AA(j)*) on gene *i* within each condition, and *β*_*IAV*_*(i)* represents the intrinsic infection effect of IAV infection. All other terms in the model are analogous to that described in M_1_. Again, the model was fit using limma, and the estimates 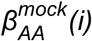 and 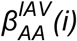 of the genetic ancestry effects were extracted across all genes, along with their corresponding p-values. Each of these estimates represents the genetic ancestry-related differential expression effects within each condition.

Genes for which the response to IAV infection is correlated with the proportion of African ancestry (i.e. popDR genes) were detected using the following model:

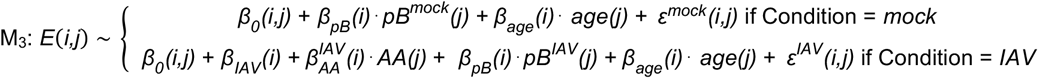

This model is similar to M_1_ (differential effect of IAV infection), in that it allows us to obtain estimates based on within-individual variability, with the difference that the IAV infection effect is no longer built in a genetic ancestry-independent manner as in model M_1_, since it is now dependent on genetic ancestry as follows: 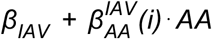 *AA*. In this context, 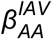 denotes the genetic ancestry-infection interaction effect induced by IAV infection, which represents variation in the response to infection that is correlated with the proportion of African ancestry.

To assess sharing of genetic ancestry effects across cell types and to increase our power to detect these effects, we applied Multivariate Adaptive Shrinkage in R (mashr v0.2.28) (Urbut et al., 2019) to the outputs of our popDE and popDR cell type-by-cell type models. mashr was applied independently to both the popDE and popDR priors, so all following methods were performed twice, once for the popDE effects and then again for the popDR effects. Effect size priors were obtained directly from limma and merged into matrices including all effect sizes across cell types, only keeping those genes detected in all cell types (i.e. n × m matrices, where for popDE effects: n = 6,847 genes, m = 10 conditions [mock- and IAV-infected popDE effects for each of the 5 main cell types], and for popDR effects: n = 6,847 genes, m = 5 conditions [popDR effects for each of the 5 main cell types]). Standard errors of the effect size priors were calculated per gene by multiplying the square root of the posterior variance (s2.post) of each gene by the unscaled standard deviation for the effect size of interest for that gene (stdev.unscaled) estimated by limma, and these values were similarly formatted into matrices as described above. To account for correlations among measurements across conditions in our data, we used the estimate_null_correlation_simple function implemented in mashr to specify a correlation matrix prior to fitting the mash model. We included both the canonical covariance matrices provided by default in mashr and data-driven covariance matrices (defined as the top 5 PCs from a PCA performed on the significant (lfsr < 0.05) signals detected in the condition-by-condition model results) learned from our data in the mash model fit. For both popDE and popDR effects, the mash model was fit to all tests using the mash function. Posterior summaries of the effect sizes, standard deviations, and measures of significance were extracted. We used the estimated local false sign rate (lfsr) to assess significance of our posterior popDE and popDR effects and considered genes significantly population differentially-expressed or differentially-responsive if the lfsr of the posterior mean was < 0.10.

### eQTL mapping and integration with mashr

eQTL mapping was performed independently in each cell type against the sets of genes retained after lowly-expressed gene filtering (n genes: CD4^+^ T cells = 9,291, CD8^+^ T cells = 9,960, B cells = 9,335, monocytes = 10,424, NK cells = 7,109, PBMCs = 10,430). A linear regression model was used to examine associations between SNP genotypes and expression levels, in which expression levels were regressed against genotype. Input expression matrices were quantile-normalized within condition prior to running the association. Mock-exposed and IAV-infected eQTL were mapped separately across all cell types. All regressions were performed using the R package MatrixEQTL (v2.3) (Shabalin, 2012). Only SNPs with a minor allele frequency > 5% across all individuals were tested, and SNPs with > 10% of missing data or deviating from Hardy-Weinberg equilibrium at p < 10^−5^ were excluded (--maf 0.05 --geno 0.10 --hwe 0.00001 PLINK v1.9 filters, www.cog-genomics.org/plink/1.9/) (Chang et al., 2015). In total, 6,305,923 SNPs passed our quality-control filters. Local associations (i.e. putative *cis*-eQTL) were tested against all SNPs located within the gene body or 100kb upstream and downstream of the transcription start site (TSS) and transcription end site (TES) for each gene tested. We recorded the minimum p-value (i.e. the strongest association) observed for each gene, which we used as statistical evidence for the presence of at least one eQTL for that gene. To estimate an FDR, we permuted the genotype data ten times, re-performed the linear regressions, and recorded the minimum p-values for the gene for each permutation. These sets of minimum p-values were used as an empirical null distribution and FDRs were calculated using the method described in the section “Modeling global infection effects”.

Power to detect *cis*-eQTL can be increased by accounting for unmeasured surrogate confounders. To identify these confounders, we first performed PCA on a correlation matrix based on gene expression for mock and IAV-infected samples. Subsequently, up to 20 principal components (PCs) were regressed out prior to performing the association analysis for each gene. A specific number of PCs to regress in each condition and cell type, corresponding to the number of PCs that empirically led to the detection of the largest number of eQTL in each condition, was then chosen from these results. The exact number of PCs regressed in each of the analyses can be found in Table S11. Of note, while PC corrections increase our power to detect eQTL, they do not affect the underlying structure of the expression data.

Mapping was performed combining both EA and AA individuals to increase power. To avoid spurious associations resulting from population structure, the first two eigenvectors obtained from a PCA on the genotype data using SNPRelate (v1.20.1, gdsfmt v1.22.0) (Zheng et al., 2012) were included in the Matrix eQTL model as well. Other covariates included in the linear model were the following: the condition and cell type-specific bimodal proportion and age (mean-centered, scaled), with two additional covariates included when mapping eQTL using the PBMC expression data, corresponding to the first 2 PCs from the cell type composition PCA described in “Modeling global infection effects”.

Our ability to detect eQTL was highly dependent on the number of cells identified in each cell type cluster (correlation between the total number of cells recovered per cell type across all individuals/conditions versus the number of significant eQTL (FDR < 0.10) detected: adj R^2^ = 0.983, p = 1×10^−8^). To gain power to detect *cis*-eQTL effects using sharing information across cell types, we again implemented mashr (Urbut et al., 2019). Out of necessity of the method, we only considered shared genes that were tested across all cell types (n = 6,573). For each of these genes, we chose a single, top *cis*-SNP, defined as the SNP with the lowest FDR across all cell types (n = 5) and conditions (n = 2), to input into mashr, yielding a total of 6,573 gene-SNP pairs. We extracted the prior effect sizes (betas) and computed the standard errors (SEs) of these betas (defined as the beta divided by the t-statistic) from the Matrix eQTL outputs for each gene-SNP pair across cell types and conditions. We defined a set of “strong” tests (i.e. the 6,573 top gene-SNP associations) as well as a set of random tests, including both null and non-null tests, which we obtained from randomly sampling 200,000 rows of a matrix containing all gene-SNP pairs tested by Matrix eQTL merged across conditions. Our mashr workflow was as follows: i) the correlation structure among the null tests was learned using the random test subset, ii) the data-driven covariance matrices were learned using the strong test subset, iii) the mash model was fit to the random test subset using canonical and data-driven covariance matrices, with two additional “infection” covariance matrices (i.e. one matrix capturing shared effects in only the mock-exposed samples and another matrix capturing shared effects in only the IAV-infected samples), and iv) the posterior summaries were computed for the strong test subset. We used the estimated local false sign rate (lfsr) to assess significance of our posterior eQTL effects and considered a gene-SNP pair to have a significant eQTL effect if the lfsr of the posterior mean was < 0.10, which we defined as an eGene.

### Identification of condition-specific popDE genes and eGenes

Within each cell type, we considered either popDE genes or eGenes as condition-specific (i.e. only showing an effect in either the mock or IAV infection condition) if they had an lfsr < 0.10 in only one condition. Here, we assume that the risk of identifying a true effect in both mock and IAV-infected cells (i.e. a shared popDE gene/eGene) as falsely condition-specific due to lack of power is low, specifically because we employed the multivariate adaptive shrinkage framework, which draws information across conditions to make better-informed posterior estimates about the sharing of effects, so we do not expect to see many posterior effects called as “condition-specific” when, in fact, they are not.

### Enrichment of eGenes within popDE genes

We tested for an enrichment of eGenes among the genes identified as popDE genes within each cell type and condition. For each cell type, condition pair, we created two vectors: i) a popDE gene vector, where significant popDE genes (lfsr < 0.1) were coded as a 1 and non-significant popDE genes were coded as a 0, and ii) an eGene vector, where significant eGenes (lfsr < 0.10) were coded as a 1 and non-significant eGenes were coded as a 0. The logistic regression was performed on the popDE gene and eGene vectors using glm in R, where the eGene vector was used as the predictor variable and the popDE gene vector was used as the response variable (popDE[0,1] ∼ eGene[0,1]). The odds ratios output by glm were converted to log2 fold enrichments with a 95% confidence interval (plotted along the x-axis in Fig. 3C).

### Calculation of predicted and observed population differences in expression

We estimated the predicted *cis*-genetic population differences in gene expression using a method in which we first computed the predicted expression of each gene considering only the posterior effect size of the top *cis* SNP for that gene and an individual’s genotype dosage (a vector of 0, 1, or 2), where, for gene *i*, individual *j*:

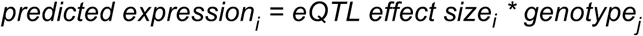

We then modeled these predicted expression values using a model analogous to that of M_2_ (model evaluating the popDE effects, “Modeling genetic ancestry effects and integration with mashr”) to obtain the predicted genetic ancestry effects (plotted on the x-axis for genetically-driven popDE genes in Fig. 3D). The observed population differences in expression were taken directly from the post-mash beta estimates of M_2_ (plotted on the y-axis for genetically-driven popDE genes in Fig. 3D).

### Modeling the effect of *cis*-regression on the observed population differences in expression

To assess the impact of *cis*-regression on population-associated expression differences, we used two models evaluating the effect of continuous genetic ancestry (African ancestry proportion) on gene expression: i) a model analogous to M_2_ (model evaluating the popDE effects, “Modeling genetic ancestry effects and integration with mashr”), and ii) a model in which, for each gene, the top *cis* SNP for that gene was regressed by including the genotype dosage for that SNP across individuals as a covariate in the model. The models were fit using limma and mashr was applied (as described in the section “Modeling genetic ancestry effects and integration with mashr”) to the prior effect sizes and standard errors derived from both models. The mashr posterior summaries were used to directly obtain the observed population differences in expression for each gene.

For each significantly enriched GO term (FDR < 0.01) identified in Fig. 3E (see “Enrichment analyses” section below), we calculated summaries of the observed population difference in expression among the genes that belong to each term that are also popDE genes with evidence of an eQTL in at least one cell type. To do this, for each cell type for each term, we collected the observed population differences among these term-specific genetically-driven popDE genes and calculated the median and standard error (SE) for these values (plotted on the x-axis in Fig. 3G). This was performed for both the observed (“real”) model outputs (model i) as well as the *cis*-regressed model outputs (model ii). For each cell type, we obtained a p-value for the real effects using a permutation method. To obtain a null distribution, we performed 1,000 permutations where, for each iteration, we: 1) sampled the same number of observed term-specific, genetically-driven popDE genes for that cell type from a background set of all genetically-driven popDE for that cell type, 2) obtained the population differences in expression among these genes, and 3) calculated the median for these null values. We then computed a one-sided, empirical p-value, where we considered the number of instances more extreme in the median null difference compared to the median observed difference in the real data given the sign of this difference (i.e. if the observed difference in the real data was < 0, we counted the number of observations in the null distribution equal to or less than the observed value, and if the observed difference in the real data was > 0, we counted the number of observations in the null distribution equal to or greater than the observed value), where p = number of instances more extreme divided by the number of permutations (n = 1000). Similarly, we obtained a p-value for the *cis*-regressed effects using the same method, except for that in steps 2 and 3, we considered the *cis*-regressed population differences as opposed to those seen in the real data. Notably, to calculate the directional p-value for the *cis*-regressed case, we used the magnitude of the median *cis*-regressed population difference but still considered the sign of the median observed population difference.

### Colocalization analysis

Specifically for the colocalization analysis, eQTL were remapped in each cell type with Matrix eQTL (Shabalin, 2012) using a 1 megabase (Mb) *cis*-window, with all other modeling parameters kept constant, to broaden our search space and increase our probability of detecting colocalized variants. We assessed colocalization between our identified eQTLs in each cell type, condition pair and 14 publicly-available GWAS summary statistics for 11 autoimmune diseases (Table S10) as previously described (Mu et al., 2020) with a few modifications. Briefly, for each trait, we identified the lead GWAS SNPs with p-values below 1×10^−5^ and defined a “locus” as a 1Mb window centered around the lead GWAS SNP. Of note, the HLA region (chr6: 25Mb-35Mb) was removed from the analysis. eGenes were defined as those with an lfsr < 0.10 from the mashr posteriors. The coloc.signals function from the coloc (v4.0.4) package in R was used to evaluate colocalization with default priors (Giambartolomei et al., 2014; Wallace, 2020). A colocalization test was only performed if the most significant SNP of an eGene fell within a GWAS locus. We defined colocalization as (PP3+PP4) > 0.5 and PP4/(PP3+PP4) > 0.8, where PP3 corresponds to the posterior probability of having two independent signals (one for the eQTL and one for the GWAS) and PP4 corresponds to the posterior probability of colocalization between the eQTL and GWAS signals.

To expand the number of colocalized genes considered for downstream analyses, we also downloaded colocalization results between harmonized bulk eQTLs in 18 immune cell types from 3 studies (DICE, DGN, and BLUEPRINT) and the same 14 autoimmune GWAS. To obtain a list of unique colocalized eGene-locus pairs, we merged colocalization results from all 18 immune cell types for each GWAS and only kept the colocalization test with the largest PP4 value for each eGene at each GWAS locus.

### Calculation of selection statistics

iHS and F_ST_ values were calculated using the 1000GP Phase 3 dataset (Auton et al., 2015) in the GRCh37/hg19 build (downloaded from ftp://ftp.1000genomes.ebi.ac.uk/vol1/ftp/release/20130502/).

### iHS

Prior to calculation, the 1000GP Phase 3 data was filtered to exclude INDELs and CNVs. Ancestral alleles were retrieved from the 6 primates EPO pipeline (version e59) (Herrero et al., 2016), and filtered 1000GP VCF files were converted to change the reference allele to the ancestral allele using bcftools (v1.9) (Li, 2011) with the fixref plugin. The program hapbin (v.1.3.0) (Maclean et al., 2015) was then used to calculate iHS values for the CEU and YRI populations using population-specific genetic maps constructed on the 1000GP OMNI dataset (ftp://ftp.1000genomes.ebi.ac.uk/vol1/ftp/technical/working/20130507_omni_recombination_rates). All downstream analyses used the standardized iHS values reported from hapbin.

### F_ST_

Prior to calculation, the 1000GP Phase 3 data was filtered to keep only biallelic SNPs. F_ST_ statistics were computed between the CEU and YRI populations using the vcftools (v0.1) (Danecek et al., 2011) flag --weir-fst-pop, where the 1000GP CEU samples were defined as population 1 and the YRI samples were defined as population 2. This method is analogous to that described in Weir and Cockerham’s 1984 paper (Weir and Cockerham, 1984).

### Enrichment of colocalized hits with popDE genes and iHS statistics

#### With popDE genes

We tested for an enrichment of popDE genes among genes with a colocalization signal across the 14 autoimmune traits included in the colocalization analysis in aggregate. Considering all colocalized signals, we collected a list of the unique genes associated with colocalization hits, corresponding to the eGenes driving the eQTL signature. Among these genes, we calculated the proportion that are also identified as popDE genes (here, a gene is considered popDE if it is called as significantly (lfsr < 0.10) popDE in at least one cell type and condition) and consider this our “observed proportion”. To obtain a null distribution, we performed 1,000 permutations where, for each iteration, we: i) sampled the same number of unique genes associated with colocalization hits from a list of all genes tested for the eQTL analysis that were shared across cell types (n = 6205), and ii) calculated the proportion of these genes that are also popDE (our “null percentage”). We calculated a p-value by evaluating the number of permutations in which the null percentage was greater than or equal to the observed percentage divided by the number of total permutations (n = 1,000).

#### With iHS statistics

We then tested for an enrichment of variants with high iHS values (defined as those with an |standardized iHS| > 95^th^ percentile of the genome-wide distribution among our tested SNPs for the population being considered) among colocalized SNPs for all of the autoimmune traits as a group. For both the 95^th^ percentile iHS calculations and our null distribution sampling approach described below, we only considered SNPs that were tested for an eQTL association in Matrix eQTL (i.e. those with a minor allele frequency > 0.05). The 95^th^ percentile |standardized iHS| cutoffs were as follows: CEU = 1.92 and YRI = 1.95. For each tested SNP, we obtained population-specific allele frequencies calculated from the 1000GP CEU or YRI individuals using the vcftools (v0.1) (Danecek et al., 2011) --freq flag. These alleles frequencies were converted to minor allele frequencies (MAFs) (i.e. if the allele frequency for a SNP was > 0.5, we subtracted it from 1), and SNPs were subsequently partitioned into 5% MAF bins (e.g. bin 1 = 0 – 5% MAF, bin 2 = 5 – 10% MAF, etc. with the rightmost interval closed).

Among the unique colocalized SNPs we identified across traits, we noticed that a subset appeared to be in high linkage disequilibrium (LD) with one another, suggesting that these SNPs likely did not represent independent colocalization signals. To account for this, we systemically identified SNPs with squared inter-variant allele count correlations (r^2^) > 0.8 using PLINK (v1.9, --r2 --ld-window-r2 0.8) (Chang et al., 2015) among all the colocalized hits with multiple tag SNPs for a single eGene. To obtain a list of independent colocalized SNPs, we included: 1) those identified as unlinked loci, and 2) only the SNP with the highest |iHS| value among each set of SNPs with an r^2^ > 0.8. We then calculated the proportion of those with an |iHS| > 95^th^ percentile and considered this our “observed percentage”. To obtain a null distribution, we used a sampling approach that mimicked both the MAF distribution and the underlying LD structure of our true data. We performed 1,000 permutations, where, for each iteration, we treated the sampling for the “independent” and “linked” SNPs separately. To create a null distribution for the “independent” SNPs, we sampled the same number of independent SNPs observed in the real data from the set of all tested SNPs in a MAF bin-matched manner, e.g. if there were 10 colocalized SNPs in MAF bin 3 in the observed data, we sampled 10 SNPs from the set of tested SNPs in MAF bin 3. To obtain a null distribution for the “linked” SNPs, we simulated the underlying LD structure of these variants where, for each eGene in the observed data with multiple tag SNPs (i.e. those SNPs with an r^2^ > 0.8), we: i) counted the number of tag SNPs for that gene and obtained the corresponding MAF bin for the SNP with the highest |iHS| value, ii) randomly sampled a set of SNPs from chromosome 1 with r^2^ > 0.8 from the MAF bin identified in i), where the number of SNPs in the set was equal to the number of tag SNPs in the observed data, and iii) picked the SNP with the highest |iHS| value among this SNP set. We then combined our simulated independent and linked SNPs and calculated the proportion of those SNPs with an |iHS| > 95^th^ percentile and considered this our “null percentage”. To calculate a p-value, we evaluated the number of permutations in which the null percentage was greater than or equal to the observed percentage divided by the number of total permutations (n = 1,000). All of the above analyses were performed twice, once with iHS values calculated within the CEU population and again with values within the YRI population.

### Enrichment analyses

Gene set enrichment analysis was performed using three independent methods, including fgsea (Korotkevich et al., 2019), GOrilla (Eden et al., 2009), and ClueGO (Bindea et al., 2009), depending on the type of data being evaluated. The enrichment program specifications and the data in which they were used to assess enrichments in are described below:

The R package fgsea (v1.10.1) was used to perform gene set enrichment analysis for the global infection effects (Fig. 1D) using the C5 gene ontology (GO) biological processes gene sets and for the popDE effects (Fig. 2D) using the H hallmark gene sets (Subramanian et al., 2005). For the infection effects, t-statistics were obtained directly from the topTable function in limma, and for the popDE effects, t-statistics were calculated from the posterior mashr outputs, where the t-statistic = posterior effect size divided by the posterior standard error for each gene. The t-statistics were then ranked, and these pre-ranked t-statistics were used to perform the enrichment using fgsea (Korotkevich et al., 2019) with the following parameters: minSize = 15, maxSize = 500, nperm = 100000. Enrichments scores (ES) and Benjamini-Hochberg adjusted p-values output by fgsea were collected for each condition and are reported in Fig. 1D and Fig. 2D for the infection effects and popDE effects, respectively.

We also used fgsea to generate the barcode plots shown in Fig. 1G to visualize where the genes in the highlighted pathways are found in the ranked specificity score list among the set of all infection differentially-expressed genes in at least one cell type. To obtain p-values for the ranked list of specificity scores, we used GOrilla (Eden et al., 2009). Notably, GOrilla relies on a statistical framework (the minimum hypergeometric score) that allows the calculation of exact p-values for observed enrichments in ranked lists of genes, taking into account multiple testing without needing to perform simulations, unlike fgsea. Because GOrilla only identifies GO terms that are significantly enriched at the top of the ranked gene list, we performed the enrichments in two ways, once with the list ranked from high to low specificity scores and again with the list ranked from low to high specificity scores. The Benjamini-Hochberg adjusted FDR q-values calculated by GOrilla for the “viral gene expression” and “response to type I interferon” terms are reported in Fig. 1G.

We performed gene set enrichment analysis for our intersection set of popDE genes and eGenes (Fig. 3E) using the ClueGO (v2.5.7) (Bindea et al., 2009) Cytoscape (v3.7.1) (Shannon et al., 2003) module in functional analysis mode, where the target set of genes was the list of popDE eGenes in the mock or IAV condition and the background set was the list of genes tested across all cell types. Specifically, we tested for the enrichment of GO terms related to biological processes (ontology source: GO_BiologicalProcess-EBI-UniProt-GOA_04.09.2018_00h00) using the following parameters: visual style = Groups, default Network Specificity, no GO Term Fusion, min. GO Tree Interval level = 3, max. GO Tree Interval level = 8, min. number of genes = 3, min. percentage of genes = 4.0, statistical test used = Enrichment/Depletion (two-sided hypergeometric test), p-value correction = Benjamini-Hochberg. For the graphical representation of the enrichment analysis, ClueGO clustering functionality was used (kappa threshold score for considering or rejecting term-to-term links set to 0.4). Only pathways with an FDR < 0.01 are reported.

**Fig. S1.**
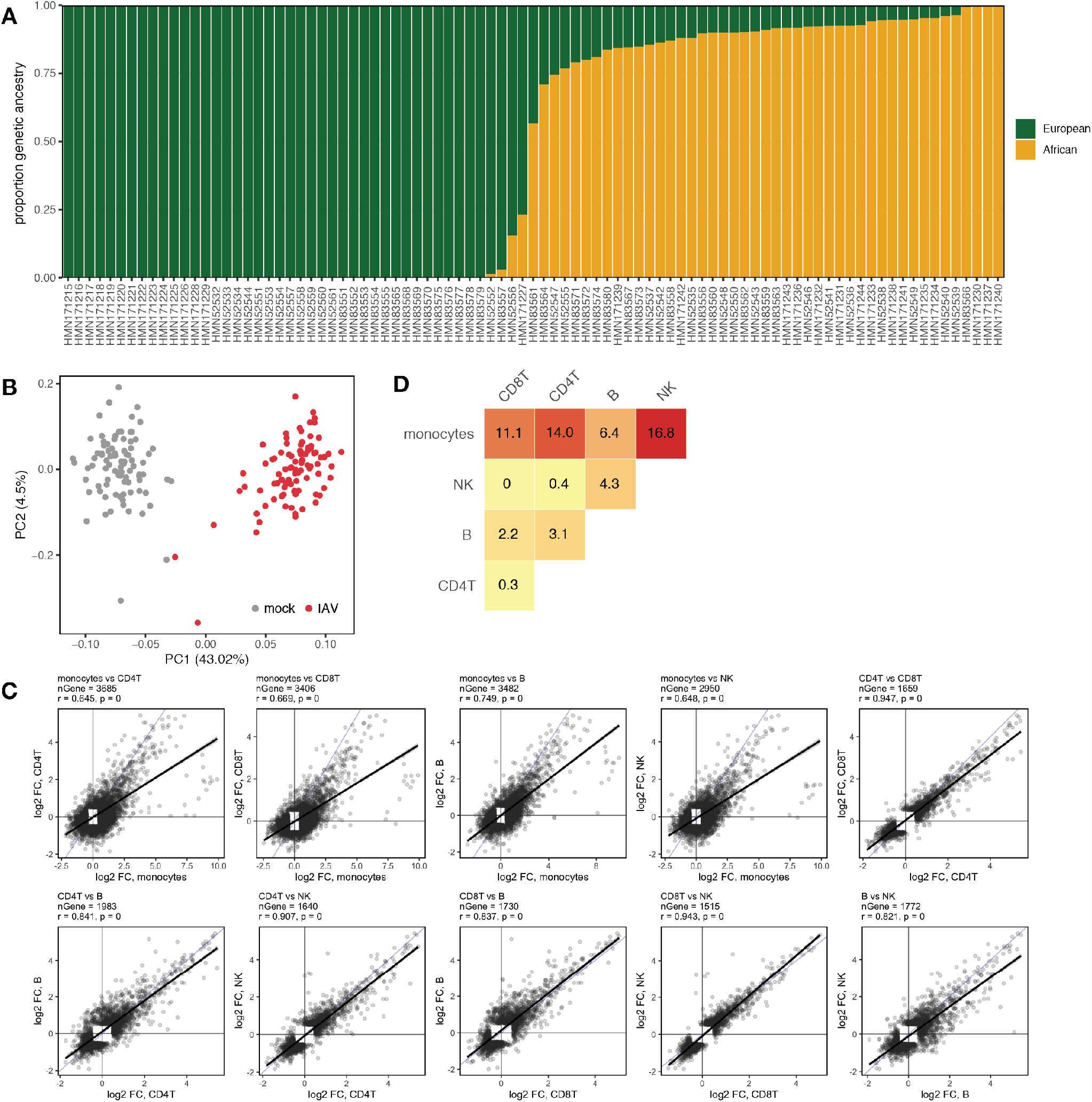
Overview of samples and global infection effects. (A) Quantitative genetic ancestry proportions partitioned into European (green) and African (yellow) components for each individual. (B) PCA decomposition of the pseudobulk PBMC expression data in mock-exposed (grey) and IAV-infected (red) samples. PC1 (percent variance explained = 43.02%) separates samples by infection status. (C) Pairwise effect size correlations across cell types among genes that are DE (logFC > 0.5, FDR < 0.05) upon IAV infection in either of the cell types being compared. (D) Pairwise comparisons of the percentage of DE genes in both cell types being compared that show discordant effect sizes.

**Fig. S2.**
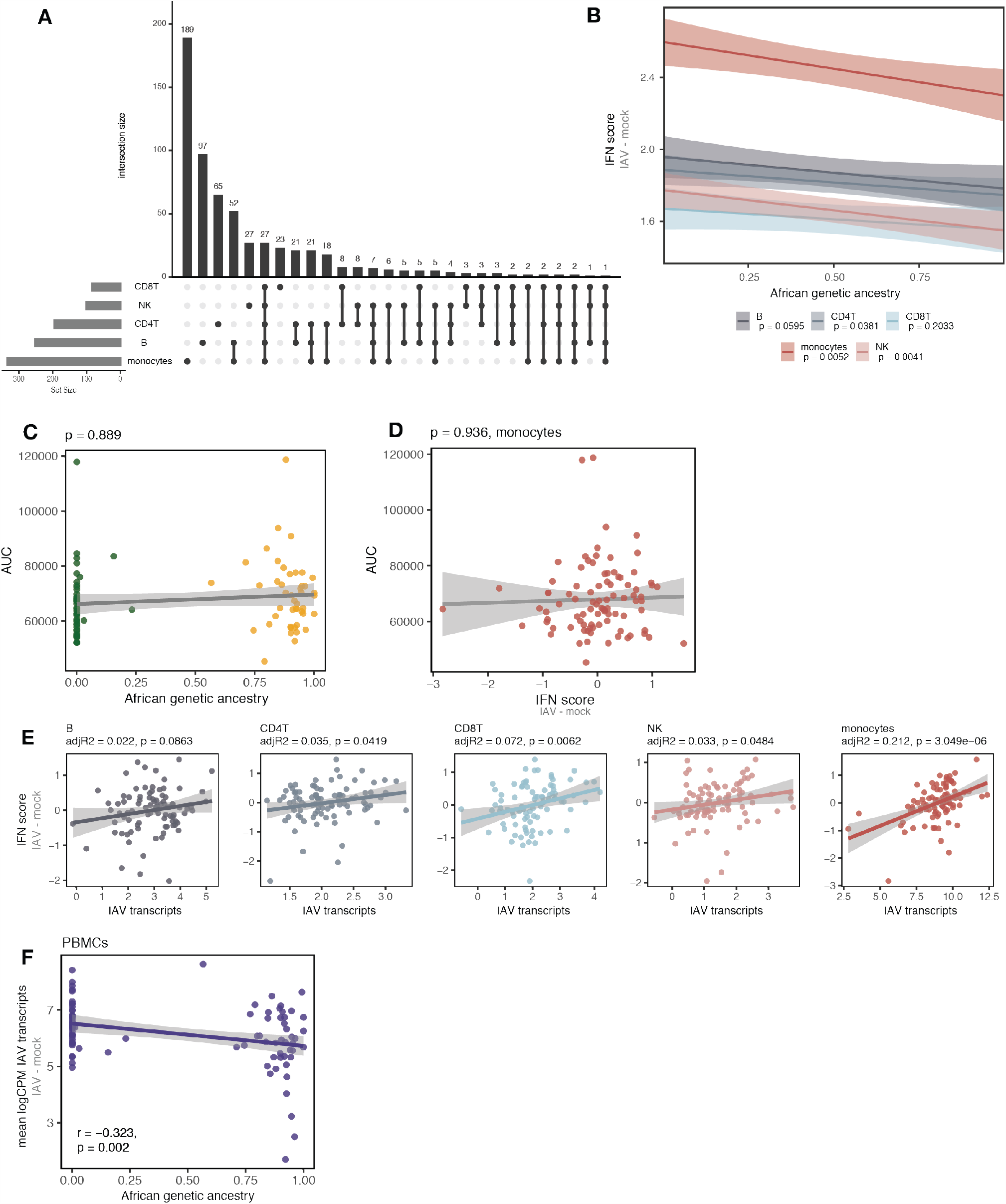
Population-associated responses to IAV infection. (A) Sharing of significant popDR genes (lfsr < 0.10) across cell types. (B) Correlation between the proportion of African genetic ancestry (x-axis) and IFN score response (y-axis) across individuals (mean Pearson’s r across cell types = −0.23, Fisher’s meta-p = 6×10^−5^). (C) Correlation between the proportion of African genetic ancestry (x-axis) and baseline levels of IAV-specific serum IgG antibodies. We quantified anti-A/Cal/04/09 antibody titers using 4-fold serial dilutions for each individual’s serum and a total of eight dilutions per sample. We then used the dilution and absorbances to generate an area under the curve (AUC; y-axis), which we used to summarize the levels of IAV (A/Cal/04/09)-specific serum IgG antibodies detected in each individual. (D) Correlation between IFN score response (x-axis) and baseline levels of IAV-specific serum IgG antibodies (“AUC” = area under the curve, y-axis). (E) Correlation between IAV transcript expression (x-axis) and the IFN response (y-axis) across individuals within each cell type. Higher IAV transcript expression is significantly associated with a stronger IFN response in CD4^+^ T cells, CD8^+^ T cells, monocytes, and NK cells (p < 0.05), with monocytes showing the strongest correlation (adj R^2^ = 0.212, p = 3.1×10^−6^). (F) African genetic ancestry is significantly negatively correlated with IAV transcript expression (Pearson’s r = −0.323, p = 0.002) in PBMCs. In (B), (C), (D), (E) and (F) lines show the best-fit slope and intercept from linear models for cell types shown.

**Fig. S3.**
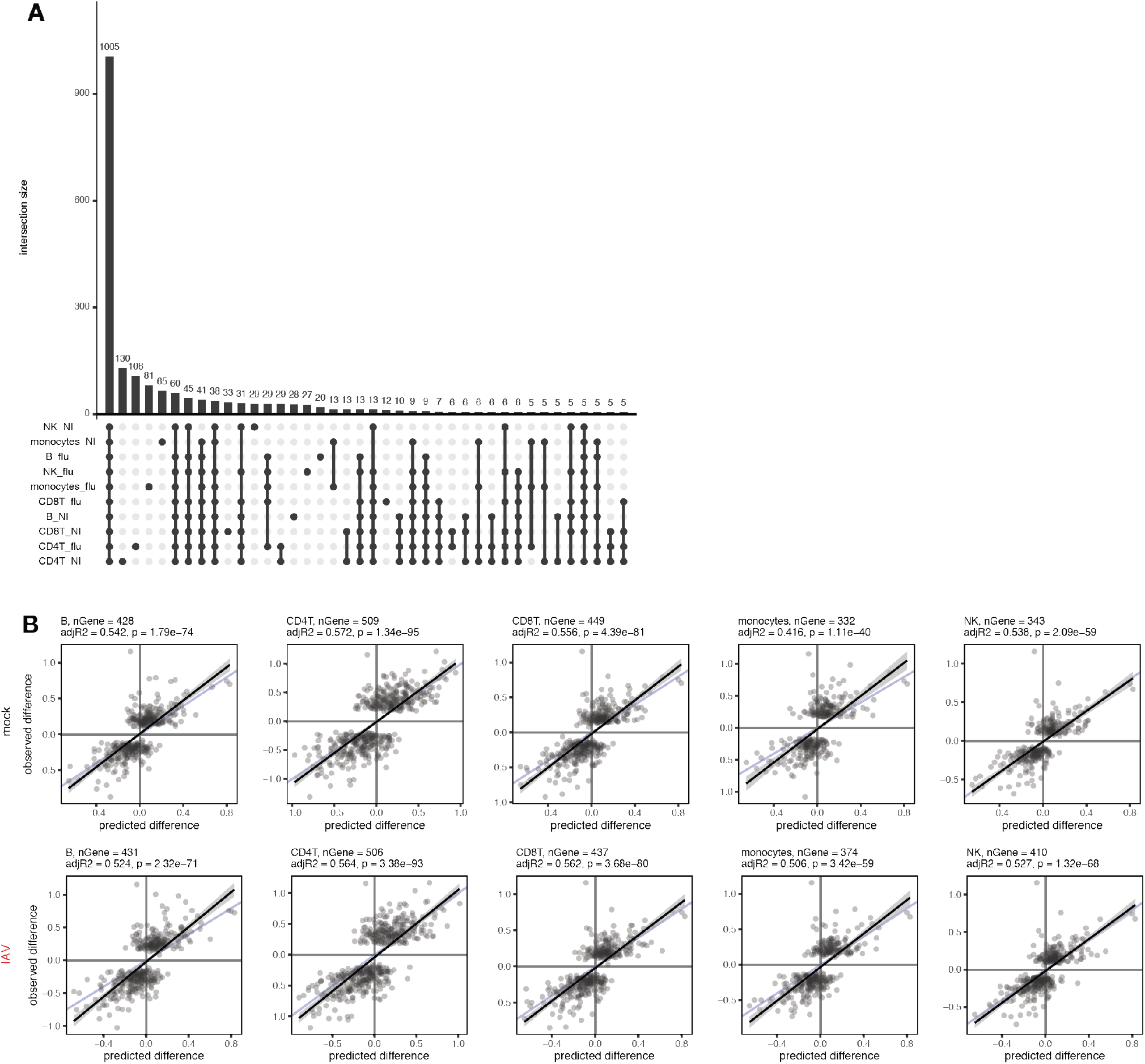
*Cis*-genetic effects regulate gene expression. (A) Sharing of significant eGenes (lfsr < 0.10) across cell types and treatment conditions. (B) Correlation of the *cis*-predicted population differences in expression (x-axis) versus the observed population differences in expression (y-axis) among popDE genes with an eQTL across all cell types in the mock-exposed condition (top) and IAV-infected condition (bottom). The black line shows the best-fit line from a linear model, and the blue line shows the identity (1:1) line.

**Fig. S4.**
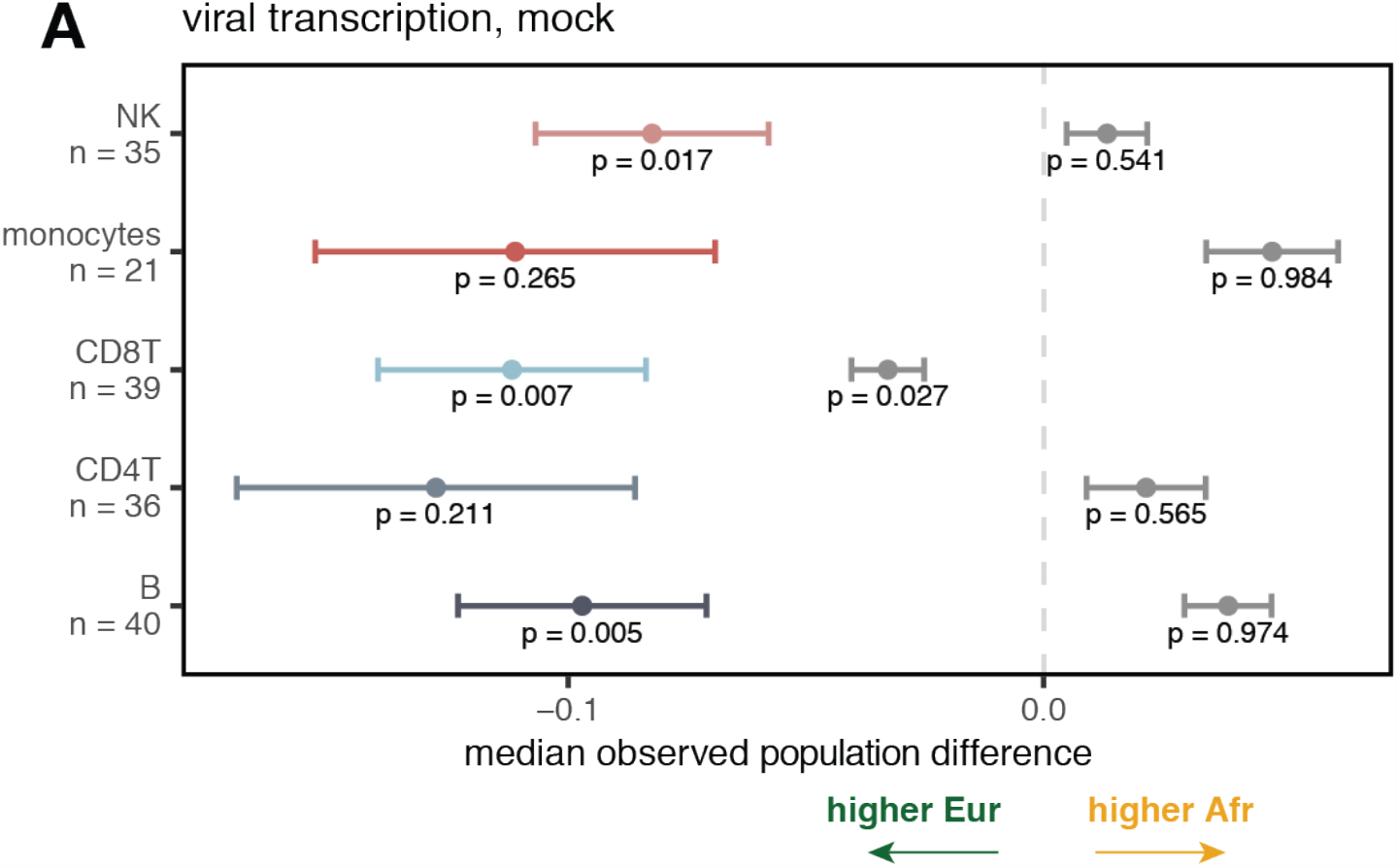
Impact of *cis*-regression on population-associated expression differences. (A) Example term showing the effect of *cis*-SNP regression. In the mock condition, EA individuals display higher expression (median observed pop. difference < 0, colored point +/- SE) of the genes belonging to the “viral transcription” term in the observed data. *Cis*-SNP regression (grey bars) reduces this effect. Points represent the median value +/- SE.

**Fig. S5.**
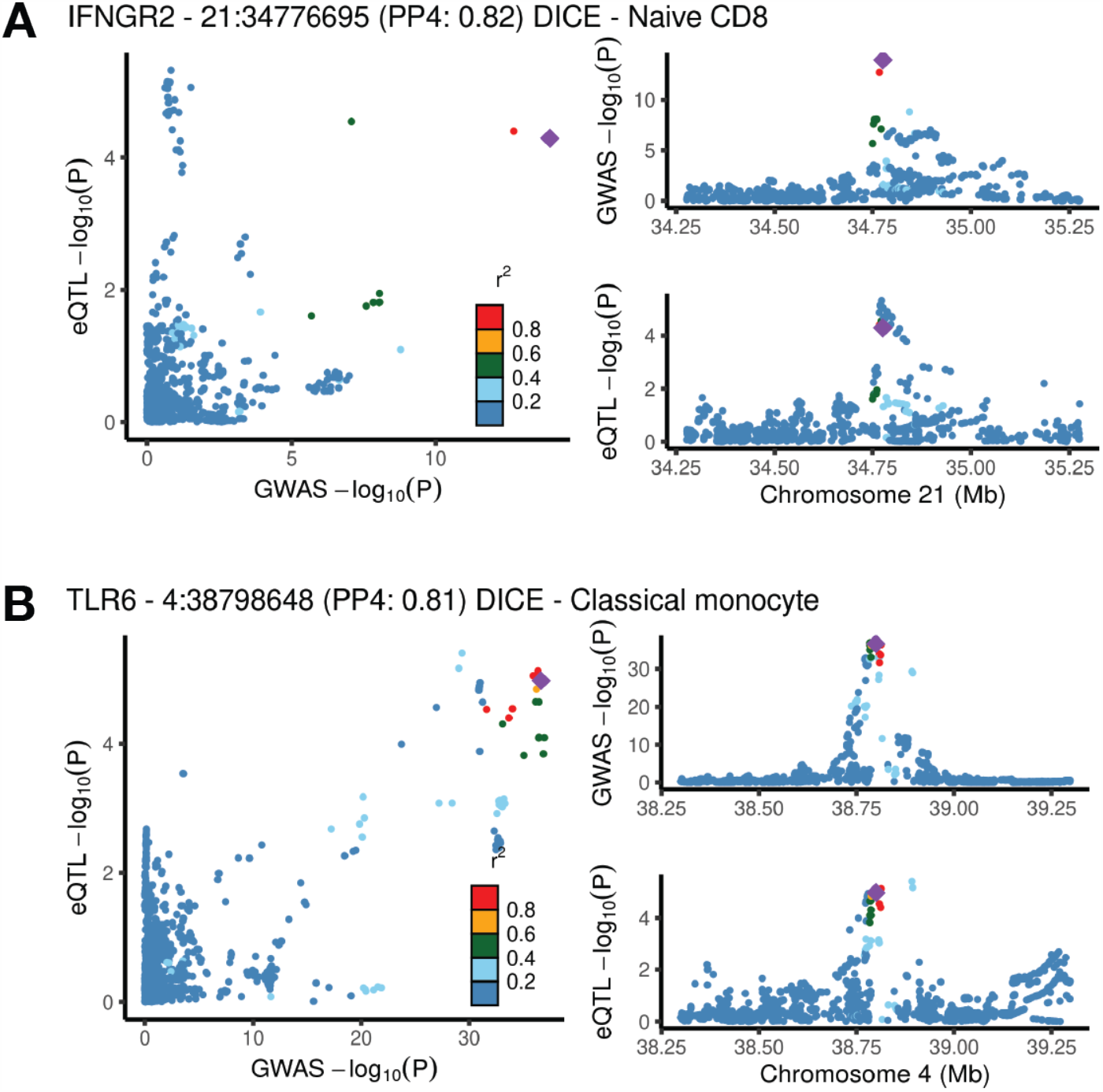
Colocalization signals. (A) *IFNGR2* colocalizes with rs2284553 in naïve CD8^+^ T cells in the Crohn’s disease GWAS (de Lange et al.). (B) *TLR6* colocalizes with rs5743618 in classical monocytes in the allergic disease GWAS. For both (A) and (B), the plot on the left shows the correlation between GWAS p-values (x-axis) and eQTL p-values (y-axis). Plots on the right show the Manhattan plots for the GWAS signal (top) and the eQTL signal (bottom).

**Table S11.** Principal components (PCs) regressed in the eQTL analysis.

| Cell type | Regressed<br>PCs (mock) | Regressed<br>PCs (IAV) | No. genes under<br>0.10 FDR (mock) | No. genes under 0.10<br>FDR (IAV) |
| --- | --- | --- | --- | --- |
| CD4<sup>+</sup> T | 1 to 4 | 1 to 2 | 1377 | 1176 |
| B | 1 to 6 | 1 to 3 | 152 | 196 |
| NK | 1 to 2 | 1 to 2 | 68 | 76 |
| monocytes | 1 to 10 | 1 to 7 | 265 | 251 |
| CD8<sup>+</sup> T | 1 to 6 | 1 to 4 | 204 | 178 |
| PBMC | 1 to 6 | 1 to 3 | 2095 | 1809 |

